# Capture of mouse and human stem cells with features of formative pluripotency

**DOI:** 10.1101/2020.09.04.283218

**Authors:** Masaki Kinoshita, Michael Barber, William Mansfield, Yingzhi Cui, Daniel Spindlow, Giuliano Giuseppe Stirparo, Sabine Dietmann, Jennifer Nichols, Austin Smith

## Abstract

Pluripotent cells emerge via a naïve founder population in the blastocyst, acquire capacity for germline and soma formation, and then undergo lineage priming. Mouse embryonic stem (ES) cells and epiblast stem cells (EpiSCs) represent the initial naïve and final primed phases of pluripotency, respectively. Here we investigate the intermediate formative stage. Using minimal exposure to specification cues, we expand stem cells from formative mouse epiblast. Unlike ES cells or EpiSCs, formative stem (FS) cells respond directly to germ cell induction. They colonise chimaeras including the germline. Transcriptome analyses show retained pre-gastrulation epiblast identity. Gain of signal responsiveness and chromatin accessibility relative to ES cells reflect lineage capacitation. FS cells show distinct transcription factor dependencies from EpiSCs, relying critically on Otx2. Finally, FS cell culture conditions applied to human naïve cells or embryos support expansion of similar stem cells, consistent with a conserved attractor state on the trajectory of mammalian pluripotency.

## INTRODUCTION

Mouse embryonic stem (ES) cells are self-renewing pluripotent cell lines derived from pre-implantation embryos (Boroviak et al., 2014; Brook and Gardner, 1997; Evans and Kaufman, 1981; Martin, 1981). They undergo continuous proliferation while exhibiting the potential to enter into multi-lineage differentiation both in vitro and upon return to the embryo. ES cells can be propagated at scale in a substantially homogeneous condition termed the naïve ground state (Martello and Smith, 2014). This is achieved in defined medium by blocking pathways that induce differentiation using two small molecule inhibitors (2i) plus the cytokine leukaemia inhibitory factor (LIF) (Wray et al., 2010; Ying et al., 2008). Ground state ES cells appear suspended in a specific time window of early development (Boroviak et al., 2014; Hackett and Surani, 2014; Smith, 2017).

Upon removal of ES cells from 2i/LIF, transcription factor circuitry associated with the naïve state is extinguished (Dunn et al., 2014; Kalkan et al., 2017; Leeb et al., 2014), metabolism is reprogrammed (Kalkan et al., 2017; Zhou et al., 2012), DNA methylation levels increase genome-wide (Lee et al., 2014), the enhancer landscape is extensively remodelled (Buecker et al., 2014; Chen et al., 2018; Kalkan et al., 2019b; Yang et al., 2014), and X chromosome inactivation initiates in female cells (Sousa et al., 2018). These events occur before lineage specification (Kalkan et al., 2017; Mulas et al., 2017), as is also evident in the peri-implantation embryo (Acampora et al., 2016; Cheng et al., 2019). During this period pluripotent cells become competent for germline specification, induced either by cytokines or by forced transcription factor expression (Hayashi et al., 2011; Nakaki et al., 2013; Ohinata et al., 2009). We hypothesise that exit from naïve pluripotency heralds a formative developmental process (Kalkan and Smith, 2014; Kinoshita and Smith, 2018; Smith, 2017) whereby competence is installed for both soma and germline induction.

During peri-implantation transition naïve pluripotency transcription factors are down-regulated and ability to give rise to ES cells is lost, while transcription factors such as Otx2 and Pou3f1 are up-regulated together with *de novo* methyltransferases Dnmt3a and Dnmt3b (Acampora et al., 2016; Boroviak et al., 2014; Boroviak et al., 2015; Brook and Gardner, 1997). By E5.5 epiblast cells have gained competence for primordial germ cell induction (Ohinata et al., 2009). Cells may persist transiently in the formative stage but overall the epiblast becomes progressively regionally fated and molecularly diverse (Beddington and Robertson, 1998; Cheng et al., 2019; Lawson et al., 1991; Peng et al., 2016; Peng et al., 2019). The period of formative pluripotency in the mouse embryo is thus postulated to span from the early post-implantation epiblast after E5.0 until the egg cylinder epiblast is extensively patterned and specified by around E6.5.

Cultures termed epiblast-derived stem cells (EpiSC) are derived by exposure of embryo explants to fibroblast growth factor (FGF) and activin (Brons et al., 2007; Guo et al., 2009; Tesar et al., 2007). EpiSCs can be derived from all stages of epiblast (Kojima et al., 2014; Najm et al., 2011; Osorno et al., 2012), but invariably converge on a mid-gastrula stage phenotype. They generally display transcriptome relatedness to primed epiblast of the anterior primitive streak around E7.25 (Kojima et al., 2014; Tsakiridis et al., 2014). Thus, culture of epiblast in relatively high levels of FGF (12.5ng/ml) and activin (20ng/ml) results in a version of primed pluripotency, which is likely prescribed by these growth factor signals.

Notably, EpiSCs are refractory to primordial germ cell induction, unlike E5.5-6.5 epiblast. (Hayashi et al., 2011; Murakami et al., 2016; Ohinata et al., 2009). Naive ES cells also fail to respond to germ cell inductive stimuli, unless they are transitioned for 24-48hrs into a population termed epiblast-like cells (EpiLCs) (Hayashi et al., 2011; Nakaki et al., 2013). EpiLCs are molecularly as well as functionally distinct from both naïve ESCs and EpiSCs (Buecker et al., 2014; Hayashi et al., 2011; Kalkan et al., 2017; Smith, 2017). They are enriched in formative phase cells related to pre-streak epiblast, but are heterogeneous and persist only transiently (Hayashi et al., 2011).

Here we invested in an effort to capture and propagate stem cells representative of mouse post-implantation epiblast between E5.5-E6.0, when the formative transition is expected to be completed but epiblast cells remain mostly unspecified.

## RESULTS

### Derivation of stem cell cultures from mouse formative epiblast

We hypothesised that shielding formative epiblast cells from lineage inductive stimuli while maintaining autocrine growth and survival signals may sustain propagation without developmental progression. Nodal, FGF4 and FGF5 are broadly expressed in the early post-implantation epiblast (Haub and Goldfarb, 1991; Mesnard et al., 2006; Niswander and Martin, 1992; Varlet et al., 1997) and promote lineage capacitation in mouse ES cells (Hayashi et al., 2011; Kunath et al., 2007; Mulas et al., 2017; Stavridis et al., 2007). They are therefore candidates for supporting formative pluripotency. However, these growth factors also drive specification in the gastrula together with Wnt3 and bone morphogenetic proteins (BMPs) (Liu et al., 1999; Winnier et al., 1995).

We speculated that moderate stimulation of FGF and Nodal pathways may be sufficient to sustain a formative population in a context of Wnt inhibition and absence of BMP. However, autocrine Nodal is known to be down-regulated *in vitro* in the absence of extraembryonic tissues (Guzman-Ayala et al., 2004), therefore we added activin A (20ng/ml) as a substitute. We used the Tankyrase inhibitor XAV939 (2μM) to block canonical Wnt signalling and excluded undefined components such as feeders, serum, KSR or matrigel. E5.5 epiblasts were isolated by microdissection and plated intact in individual fibronectin-coated 4-well plates in N2B27 medium under 5% O_2_ (Fig. 1A). After 5-6 days, explants were treated with pre-warmed accutase for 5-10 seconds then gently detached, fragmented into small clumps, and seeded into fresh 4-well plates. With or without added FGF, colonies of tightly packed epithelioid cells grew up that could be passaged further and expanded into continuous cell lines (Fig. 1A and S1A). Although cultures are morphologically similar in both conditions, we detected appreciably higher expression of primitive streak markers Brachyury, FoxA2, Eomes and Gsc, in the absence of FGF (Fig S1B, C). Nodal/activin signalling is known to stimulate these genes (Brennan et al., 2001; Conlon et al., 1994; Takenaga et al., 2007). We therefore titrated activin in the absence of FGF and found that continuous cultures could still be established (Fig. 1B and S1D). In low activin (3ng/ml) plus XAV939 (A_lo_X) we obtained cell lines that could be propagated for more than 20 passages (Fig. 1B, S1D, Supplemental movie 1).

**Fig. 1.**
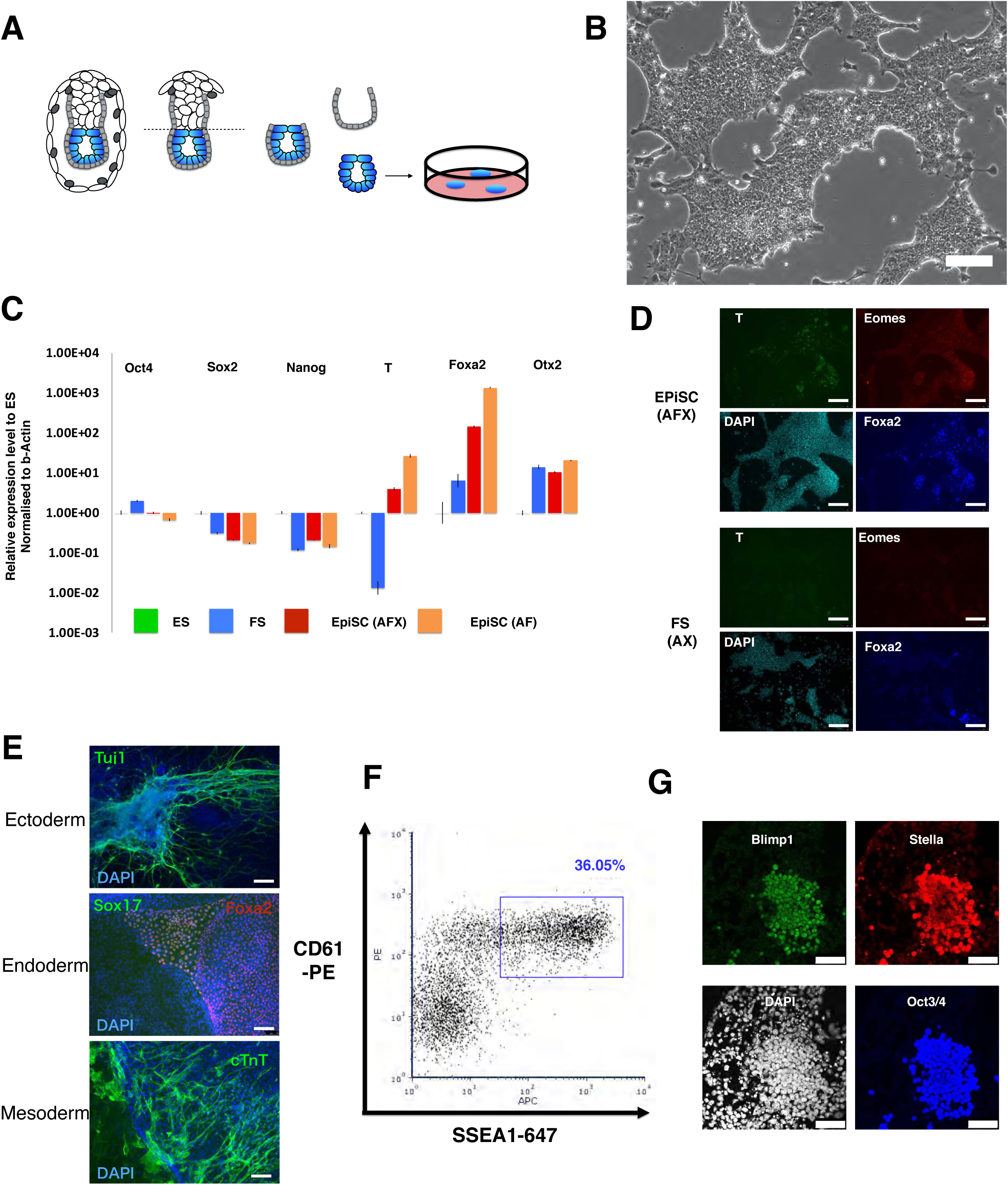
(A) Schematic drawing of cell line derivation from E5.5 epiblast. (B) Bright field image of serially passaged E5.5 epiblast derived culture. Scale bar 100µm. (C) RT-qPCR analysis of marker gene expression. Expression is relative to ES cells in 2iL (=1), normalized to beta-actin. Error bars are S.D. from technical triplicates. (D) Immunofluorescent staining of EpiSCs and epiblast-derived (FS) cells for early lineage markers, scale bars 150µm. (E) Immunostaining of embryoid body: ectoderm, Tuj1, green; endoderm, Foxa2, red, and Sox17, green; mesoderm, cTnT, green; DAPI blue. Scale bars, 150µm. (F) Flow cytometry analysis of PGCLC induction at day 4. (G) Immunostaining of day 4 PGCLC. Blimp1 in green, Stella in red, Oct4 in blue, DAPI in white. Scale bars 50 µm.

Cell lines derived in A_lo_X express *Otx2*, consistent with post-implantation identity, but show no expression of *T* and minimal *FoxA2* (Fig. 1C, D). They display similar levels of *Pou5f1 (Oct4)* mRNA to EpiSCs, slightly higher *Sox2*, and lower *Nanog*. (Fig. 1C). Upon embryoid body formation and outgrowth, we detected expression of germ layer markers, indicating multi-lineage differentiation (Fig 1E).

These observations suggest that in the absence of other stimuli, limited activation of the Nodal/activin pathway combined with autocrine stimulation of the FGF pathway may be sufficient to suspend cells in the formative phase of pluripotency.

### Stem cell propagation is facilitated by retinoic acid receptor inhibition and requires Nodal pathway activity

During establishment and expansion in A_lo_X we observed sporadic expression of neural lineage markers and differentiation predominantly into neuronal morphologies. On occasion differentiation was extensive and led to loss of cultures. We considered that retinoids might be acting as neural inductive stimuli (Bain et al., 1995; Stavridis et al., 2010). We therefore examined the effect of a pan-retinoic acid receptor inverse agonist (RARi, BMS 493; 1.0µM) (Fig. S1E). Supplementation of A_lo_X with RARi, henceforth A_lo_XR, resulted in improved derivation efficiency (Fig. S1F), reduced ectopic expression of neural specification factors Sox1 and Pax6 (Fig. S1E), and stabilisation of long-term cultures. Using A_lo_XR we established nine cell lines from embryos of two different genetic backgrounds, 129 and CD1. These lines were all passaged more than 10 times (30 generations) with no indication of crisis or senescence. Established cultures grow more slowly than EpiSCs at a similar rate to ES cells, with routine passaging every 2-3 days at a split ratio of 1/10-1/15. Chromosome counts reveal a majority of diploid cells even at later passages (Fig. S1G). Cells were routinely passaged by mild dissociation into small clumps. Survival was poor after dissociation to single cells but could be rescued by addition of ROCK inhibitor which enabled clonal expansion after genetic manipulation. We used fluorescent *in situ* hybridisation (FISH) to monitor expression of Xist in a female line. Detection of a prominent cloud of hybridisation in each nucleus (Fig. S1H) is indicative of initiation of X chromosome inactivation, distinct from naïve epiblast and consistent with observation of formative epiblast in vivo (Mak et al., 2004; Shiura and Abe, 2019).

Mouse ES cells undergo formative transition when withdrawn from 2iLIF (Hayashi et al., 2011; Kalkan et al., 2017; Mulas et al., 2017). We applied A_lo_XR during this transition and obtained continuously proliferating epithelioid cells. Cultures display variable levels of heterogeneity during the first few passages (Fig. S1I) but stabilise within 4-6 passages and subsequently expand similarly to embryo derived cell lines. To test for persistence of naive cells, we replated cultures in 2iLIF, which supports clonal propagation of ES cells at high efficiency (Kalkan et al., 2017). All cells died or differentiated within a few days, demonstrating complete extinction of ES cell identity. This finding is in marked contrast to other reports of “intermediate” pluripotent states, which readily revert to ES cells (D’Aniello et al., 2016; Neagu et al., 2020; Rathjen et al., 1999).

### Germline and somatic lineage induction in vitro

In mouse, the formative phase of pluripotency is definitively distinguished from naïve and primed phases by competence for germline specification (Hayashi et al., 2011; Ohinata et al., 2009). We examined the response of embryo-derived A_lo_XR cells to the inductive cytokine cocktail (BMP2, SCF, EGF and LIF) (Ohinata et al., 2009). We tested 8 independent lines and in each case detected the primordial germ cell surface marker phenotype CD61^+^SSEA1^+^ (Fig. 1F). This capacity was maintained even in late passage (>P30) cultures. The proportion of marker positive cells ranged up to >30% in some experiments, and was generally between 5-25%, although one line was consistently less efficient, around 1%. Two lines propagated in A_lo_X also produced CD61^+^SSEA1^+^ immunopositive cells, albeit at <10% (Fig. S1J). In contrast, 4 independent AFX EpiSC lines derived from E5.5 epiblast did not yield double positive cells (Fig. S1K). Furthermore, AFX EpiSCs adapted to culture in A_lo_XR over several passages remained unable to produce primordial germ cell-like cells (PGCLC) (Fig. S1L).

To confirm PGCLC identity, we sorted the CD61^+^SSEA1^+^ population and verified expression of a range of germ cell markers by RT-qPCR (Fig. S1M). We also observed co-expression of Oct4, Blimp1 and Stella proteins by immunostaining in both A_lo_XR and A_lo_X cultures (Fig. 1G, S1N). Collectively these features constitute recognised hallmarks of mouse PGCLC (Hayashi et al., 2011; Ohinata et al., 2005). Based on this competence we designated A_lo_X and A_lo_XR cells as formative stem (FS) cells.

We then investigated directed somatic differentiation of FS cells in comparison with EpiSCs. Inhibition of the Wnt pathway shifts the character of EpiSCs towards anterior epiblast identity and predisposes them to neuroectodermal fate (Osteil et al., 2019; Tsakiridis et al., 2014). We used the Sox1::GFP reporter (Stavridis and Smith, 2003) to quantify neural induction kinetics of FS cells and EpiSCs maintained with Wnt inhibition. At day 1 after transfer into non-supplemented N2B27 medium, more than 80% of EpiSCs are GFP positive compared with only around 25% of FS cells (Fig. 2A). By day 2, however, the GFP+ fraction approaches 80% for FS cells and by day 3 both is over 80% as for EpiSCs. We examined protein expression by immunostaining and found that FS cells lag behind EpiSCs in both down-regulation of Oct4 and up-regulation of Sox1, but by day 3 the vast majority are Oct4-negative and Sox1-positive (Fig. 2B). Thus, mouse FS cells have similar capacity to form neuroectoderm as EpiSCs, but take longer to do so.

**Fig. 2.**
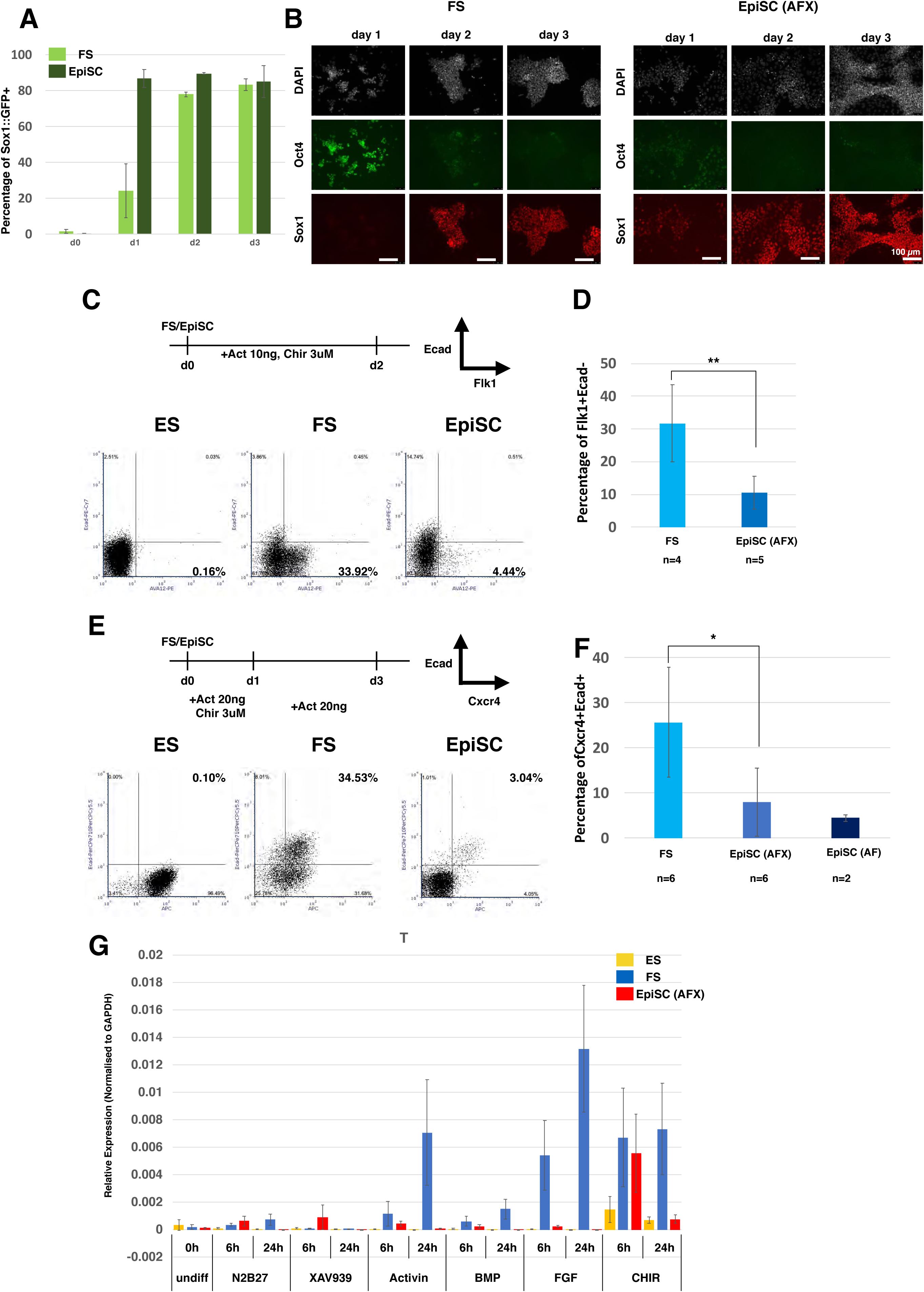
(A) Efficiency of neural differentiation assayed by Sox1::GFP. Error bars represent S.D. from 4 independent differentiation experiments. (B) Immunostaining of FS cells and EpiSCs during neural differentiation. Oct4 in green; Sox1 in red; nuclei staining with DAPI in white. Scale bars, 100 µm. (C) Protocol for lateral plate mesoderm differentiation. Representative results for ES cells, FS cells, EpiSCs. Mesoderm differentiation efficiency was assessed as the Flk1^+^Ecad^-^ fraction by flow cytometry. (D) Average efficiency of Flk1 positive fraction production from FS cells and EpiSCs. n = number of independent cell lines assayed. Error bars represent the S.D. **P<0.01. (E) Protocol for definitive endoderm differentiation. Endoderm fraction was quantified as the Cxcr4^+^Ecad^+^ fraction by flow cytometry. (F) Average proportion of Cxcr4^+^Ecad^+^ double positive fractions from differentiation of 6 FS lines, 5 EpiSC (AFX) lines (6 experiments) and 2 EpiSC (AF) lines. Error bars represent S.D., *P<0.05. (G) T expression analysed by RT-qPCR at 0, 6 and 24 hours after the indicated stimuli; 2µM XAV939, 20ng/ml activin A, 10ng/ml BMP2, 12.5ng/ml Fgf2 and 3µM CH. Relative expression is normalised to GAPDH. Error bars are S.D. from two independent cell lines and two technical replicates.

We tested primitive streak-like induction in response to activin and GSK3 inhibition (Burgold et al., 2019). We observed substantially higher induction of mesendoderm surface markers and gene expression from FS cells than from EpiSCs (Fig. S2A-C). We used flow cytometry to quantify Flk1^+^Ecad^-^ lateral mesoderm and Cxcr4^+^Ecad^+^ definitive endoderm. We observed no induction of either lineage directly from ground state ES cells and relatively poor induction from EpiSCs compared to FS cells (Fig. 2C and 2E). We examined several independent FS lines and EpiSC lines. Induction of mesoderm was on average three-fold more efficient from FS cells (Fig. 2D), and of endoderm four-fold higher (Fig. 2F).

To investigate the basis of differential propensity for primitive streak induction we examined the response of ESCs, FS cells and EpiSCs to signalling pathways active during gastrulation. Ground state ESCs did not up-regulate *T* in response to any stimulus tested with the exception of very low induction by the GSK3 inhibitor CH. EpiSCs also failed to show any appreciable response, apart from induction by CH at 6hrs that was not maintained at 24hrs. In contrast, FS cells showed sustained up-regulation of *T* upon treatment with activin, FGF, CH, or, to a lesser extent, BMP (Fig. 2G). Notably, FGF as low as 1 ng/ml induced T and FoxA2 expression in FS cells (Fig. S2D)

Thus, FS cells show rapid and efficient responsiveness to primitive streak inductive cues but require 48 hours to elaborate neural specification. These behaviours are distinct from EpiSCs, and consistent with a developmental stage of E5.5-6.0 epiblast.

### Chimaera colonisation

AF EpiSCs do not normally make contributions to blastocyst injection chimaeras, unless they have been genetically modified to enhance ICM integration or survival (Masaki et al., 2016; Ohtsuka et al., 2012; Tesar et al., 2007). We tested AFX EpiSCs derived from E5.5 epiblast and observed no chimaeras after blastocyst injection of three lines and transfer of 95 embryos. We reasoned that FS cells may have higher probability of persisting from injection into the E3.5 blastocyst until developmentally stage matched early post-implantation epiblast emerges. We engineered embryo-derived FS cells to express mKO2 or GFP fluorescent reporters. After blastocyst injection and uterine transfer we detected reporter expression in multiple mid-gestation foetuses from three different FS cell lines (Fig. 3A, Fig. S3A-E). Contributions are low to moderate compared with typical ESC chimaeras and tend to be patchy rather than evenly dispersed. Nonetheless, colonisation may be spread over multiple tissue types, including Sox2 positive putative migratory primordial germ cells at E9.5 (Fig. 3B). We examined genital ridge contribution at E12.5 and detected mKO2 reporter positive Oct4^+^ Mvh^+^ primordial germ cells (Fig. 3C, S3F, G). We also observed contributions to 3 newborn pups by fluorescence imaging. Two animals developed to adulthood but the other was euthanised at P21 due to malocclusion. Post-mortem tissue inspection revealed contributions to brain, bone, skin, heart, lung and gut (Fig. 3D).

**Fig. 3.**
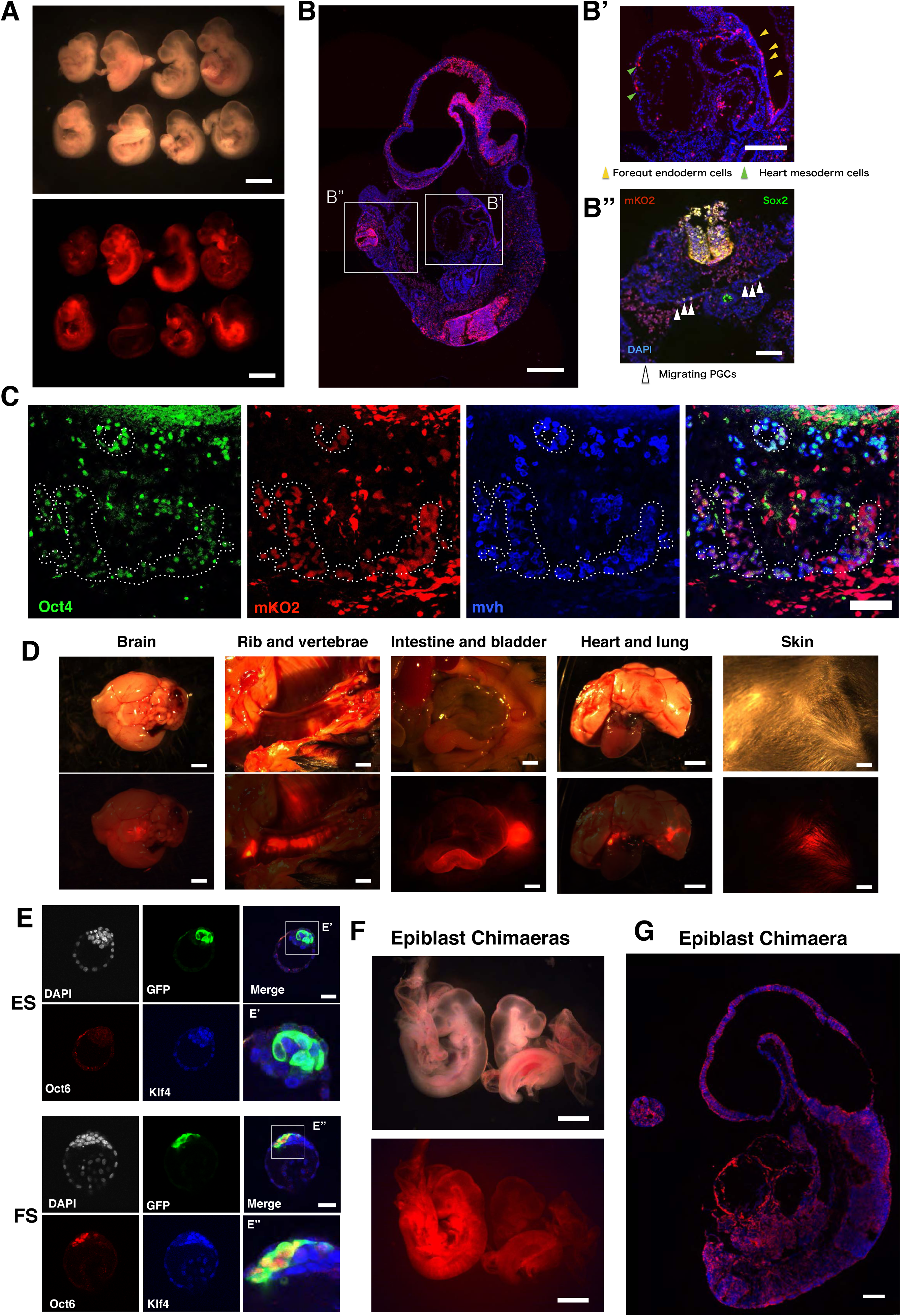
(A) Bright field and fluorescent images of E9.5 chimaeric embryos generated by blastocyst injection of mKO2 reporter FS cells. Scale bar is 1mm. (B) Sagittal section from one chimaera, mKO2 and DAPI stained. Inset B’, mKO2 positive cells in foregut endoderm (yellow arrowheads) and cardiac mesoderm cells (green arrowheads). Inset B’’ (rotated 90^0^), additionally showing Sox2 immunostaining with putative migrating PGCs in the hindgut region indicated by white arrowheads. Scale bars: (B) 200µm; (B’, B”) 100µm. (C) mKO2 positive cells expressing Oct4 and Mvh PGC markers in E12.5 gonad. Triple positive cells are highlighted with dashed circles. Scare bar, 75µm. (D) Contributions of mKO2 positive FS cells in post-natal chimaeras. Fluorescent images of brain, rib and vertebrae and heart and lung were overlaid with 20 % opacity bright field image. Scale bars, 2 mm. (E) Blastocyst chimaeras generated with GFP reporter ES cells or FS cells and cultured for 24 hours. ES cells are Klf4^+^Oct6^-^ (n=11) (E’) whereas FS cells are Klf4^-^Oct6^+^ (E’’) (n=15). Scale bars, 40 µm. (F) E9.5 Chimaeras obtained from blastocyst injection of donor membrane tdTomato expressing E5.5 epiblast cells. Scale bars, 500 µm. (G) Section from the left embryo in Panel F stained with anti-RFP to visualise tdTomato. DAPI staining is in blue. Scale bar, 200 µm.

To examine whether chimaera colonisation might entail reversion of FS cells to naïve status in the blastocyst environment we inspected embryos 24 hours after injection. We found that the labelled donor cells were appropriately localised within the ICM. However, immunostaining showed that in contrast to host naïve epiblast or introduced ESCs, FS cells did not express the naïve pluripotency specific transcription factor Klf4 and retained the formative marker Oct6 (Fig. 3E). Therefore, FS cells maintain formative identity within the blastocyst environment.

Chimaera formation by FS cells derived from post-implantation epiblast challenges the conclusion from classic embryo-embryo chimaera studies that epiblast cells lose colonisation ability entirely by E5.5 (Gardner and Brook, 1997; Gardner et al., 1985). We revisited those experiments using a fluorescent reporter to allow sensitive detection of donor epiblast cell contributions. We also added Rho-associated kinase inhibitor to improve viability of isolated epiblast cells. We dissected epiblasts from cavitated E5.5 and pre-streak E6.0-6.25 transgenic embryos that constitutively express membrane-bound tdTomato (mTmG). The epiblasts were dissociated with Accutase and 10 single epiblast cells were injected per blastocyst. We detected tdTomato positive cells in 11 out of 91 embryos recovered at E9.5 (Fig. 3F, G, S3H-S3L). Contributions were typically sparse and interestingly were most frequent in yolk sac mesoderm and amnion. In three chimaeras, however, colonization was widespread in the embryo proper (Fig. 3F, G, S3H). We did not detect any contribution from early streak stage epiblast donor cells (Fig. S3L).

These observations establish that embryonic formative epiblast cells can contribute to blastocyst chimaeras, although with lower efficiency than ICM or ES cells.

### Transcriptome relatedness to pre-streak epiblast

For global evaluation of cellular identity we performed RNA-seq. We first compared FS cells with ground state ES cells and EpiSCs cultured in AF or AFX. Principal component analysis (PCA) separates ES cells on PC1 while PC2 resolves the two types of EpiSCs and FS cells (Fig. 4A). Differential expression analysis (Log_2_ fold change > 1.4, adjusted P value < 0.05) identified 531 and 266 genes up-regulated and 941 and 168 genes down-regulated in FS cells compared with the AF and AFX EpiSCs respectively (Fig S4A ad S4B). GO term enrichment analysis highlights “cell adhesion” in FS cells in contrast to terms related to gastrulation and development in EpiSCs (Fig. S4A and S4B). We identified 328 genes that are up-regulated in FS cells compared with ES cells or either class of EpiSC (Fig 4B). GO term analysis shows terms related to “ion transport” and “cell adhesion” (Fig. 4C).

**Fig. 4.**
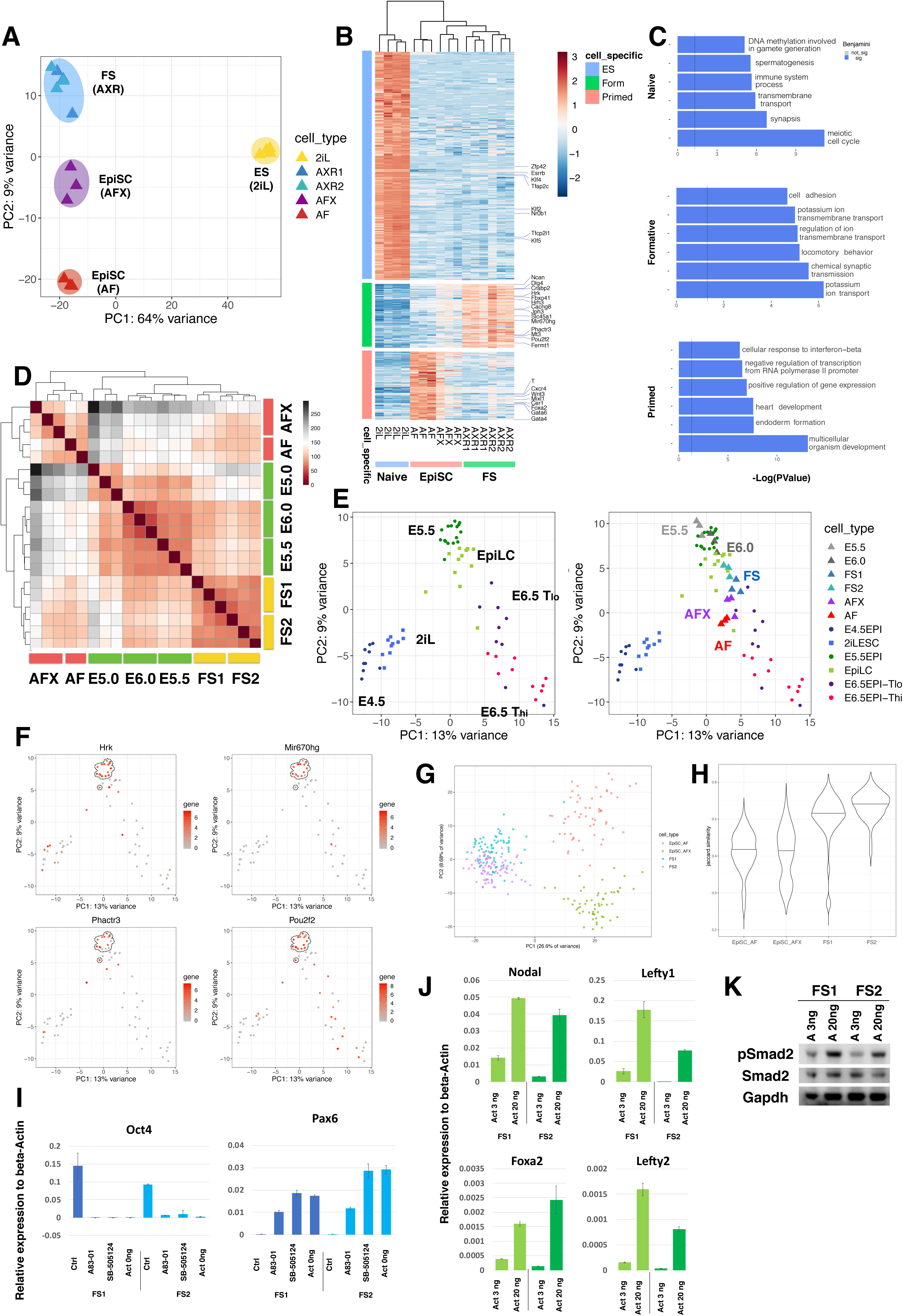
(A) PCA with all genes for ES cells, FS cells and EpiSCs (AFX and AF). (B) Heatmap clustering of naïve, formative and primed enriched genes. (C) GO term analyses based on the genes identified in (B). X-axis is -Log(P-Value). Top 6 significant terms are shown (Benjamini value<0.05). (D) Heatmap comparison of FS cells and AFX and AF EpiSCs with E5.0, E5.5 and E6.0 epiblast cells. (E) PCA analysis with mouse single cell data during pre-implantation and gastrulation stage embryo (left) (Nakamura et al 2016). Samples from (D) were projected onto the single cell PCA (right). (F) Gene expression pattern of selected FS cell enriched genes identified in (B) coloured on PCA. E5.5 epiblast cells are highlighted by dashed circle. (G) PCA plot of scRNA-seq data from two FS cell lines and one AFX and one AF EpiSC line. 2,000 most abundant genes were used. (H) Violin plot of Jaccard index analysis of 2,000 most abundant genes shows higher correlation between FS cells than EpiSCs. (I) RT-qPCR analysis of FS cells in AloXR (Ctrl) or with addition of 1µM A83-01, 5 µM SB5124 treated or withdrawal of activin for 2 days. Relative expression to beta-actin. Error bars are S.D. from technical duplicates. (J) RT-qPCR analysis of FS cells cultured in low (3ng/ml) and high (20ng/ml) activin for two days. Relative expression to beta-actin. Error bars are SD from technical duplicates. (K) Western blot analysis of phospho-Smad2 protein. Cells were passaged once with low (3ng/ml) or high (20ng/ml) activin A concentration before collecting protein.

We then used a low cell number bulk RNA-seq protocol with a comprehensive read depth (Boroviak et al., 2015) for direct comparison of FS cells with dissected pre-cavitation (E5.0), early cavitation (E5.5), and pre-streak (E6.0) epiblast. Unsupervised hierarchical clustering shows that FS cells are most related to E5.5 and E6.0 epiblast, with lower correlation to the pre-cavitation stage (Fig. 4D). EpiSCs, both AF and AFX, form a separate cluster, less related to these pre-gastrula epiblast stages. We identified 953 differentially expressed genes between FS cells and EpiSCs. This gene set clusters published embryo and EpiLC single cell data (Nakamura et al., 2016) according to developmental trajectory (Fig. 4E). When projected onto this scRNA-seq PCA, bulk RNAseq E5.5 and E6.0 epiblast align with E5.5 and EpiLC samples as expected (Fig. 4E). FS cells sit between E5.5 and E6.5 T_Lo_, and overlap with EpiLCs, whereas EpiSCs position with the E6.5 cells. We inspected several of the FS cell specific genes identified in vitro (Fig 4B) and detected dynamic expression in the embryo with enrichment at E5.5 (Fig. 4F, S4C).

We also performed single cell analysis using the Smart2-seq method (Picelli et al., 2014). Cells with fewer than 3M reads were removed from further analysis, leaving 326 cells that passed the threshold. FS cells from two independent lines formed a single cluster in the PCA plot (Fig. 4G), separated from EpiSCs on PC1. Notably there was no overlap between EpiSCs and FS cells. PC2 separates AF and AFX EpiSCs. Measurement of gene expression correlation by Jaccard index confirmed that FS cells are more homogeneous than either class of EpiSC (Fig. 4H).

Collectively these analyses indicate that FS cells capture features of pre-streak epiblast and are closely related to EpiLCs, but less related to EpiSCs.

### Growth factor requirements for FS cell propagation

We evaluated Nodal, FGF and Wnt family representation in the FS cell transcriptome data (Fig. S4D-F) as potential autocrine stimuli of self-renewal or differentiation. As expected for formative cells, we found robust expression of FGF5 but also detected several other FGFs at lower levels. Interestingly, FGF8 implicated in primitive streak formation, is lowly expressed compared with EpiSCs. FS cells express both FGFR1 and FGFR2 (Fig. S4D). We tested whether FS cell cultures are dependent on FGF signalling by adding specific inhibitors of the FGF receptor (PD173074, 0.1µM) or downstream MEK1/2 (PD0325901,1µM). Both inhibitors caused rapid collapse of FS cell cultures. We conclude that endogenous low-level expression of FGFs supports self-renewal, without inducing the primitive streak-associated gene expression associated with exposure to exogenous FGF (Fig. 2G, S2D).

FS cells express nodal/activin receptors but interestingly present lower levels of the co-receptor Tdgf1 and of Nodal than either ES cells or EpiSCs (Fig. S4E). We investigated further the requirement for nodal pathway stimulation. Addition of receptor inhibitors (A83-01 or SB505124) resulted in extensive cell death and differentiation with loss of Oct4 and up-regulation of Pax6 (Fig. 4I and S4G). Withdrawal of activin also led to reduced viability and increased differentiation, indicating that autocrine effects do not provide sufficient pathway activation. In FS cell medium activin is added at only 3ng/ml, however, compared with 20ng/ml typically used for feeder-free culture of EpiSCs. Dosage sensitivity is a well-known feature of nodal signalling in the mouse embryo (Robertson, 2014). We observed markedly less induction of nodal pathway targets in FS cells at 3ng/ml compared to 20ng/ml activin (Fig. 4J). Furthermore, immunoblotting indicated lower steady state levels of phospho-Smad2 in cells passaged in 3ng/ml activin (Fig 4K). These observations are consistent with a dose-dependent response to nodal/activin stimulation, whereby low signal sustains formative cells and high signal promotes primitive streak specification.

Finally, the expression of Fzd receptors and low levels of some Wnts may underlie the requirement for inhibition of Wnt signalling to fully suppress differentiation (Fig. S4F). Consistent with this interpretation we observed that the porcupine inhibitor IWP2 could substitute for XAV939 during FS cell maintenance.

Thus, FS cells are maintained by FGF and nodal/activin but are poised to respond to increased levels of either signal or of canonical Wnt by entering into mesendoderm differentiation.

### Chromatin accessibility in formative stem cells

We employed the assay for transposase accessible chromatin coupled to deep sequencing (ATAC-seq) (Buenrostro et al., 2013) to survey open chromatin in FS cells. Independent FS cell samples were well correlated (Fig. 5A). We classified sites that exhibit differential accessibility between ES, FS and EpiSCs based on a fold-change enrichment greater than two (p-value<0.05). A major re-organisation is evident between naïve and formative cells, with 3742 sites closing, 4259 opening and only 207 shared open sites (Fig. 5B,C). In contrast, between formative and primed cells, a majority of open sites are shared (3588), while just over 1000 become more accessible and a similar number close. We detected 826 peaks specifically enriched in FS cells compared to either ES cells or EpiSCs (Fig. 5B, C). These FS cell-specific open chromatin regions are also accessible in transient EpiLCs (Fig. 5C and 5D) but no significant GO terms are enriched (Fig. S5A).

**Fig. 5.**
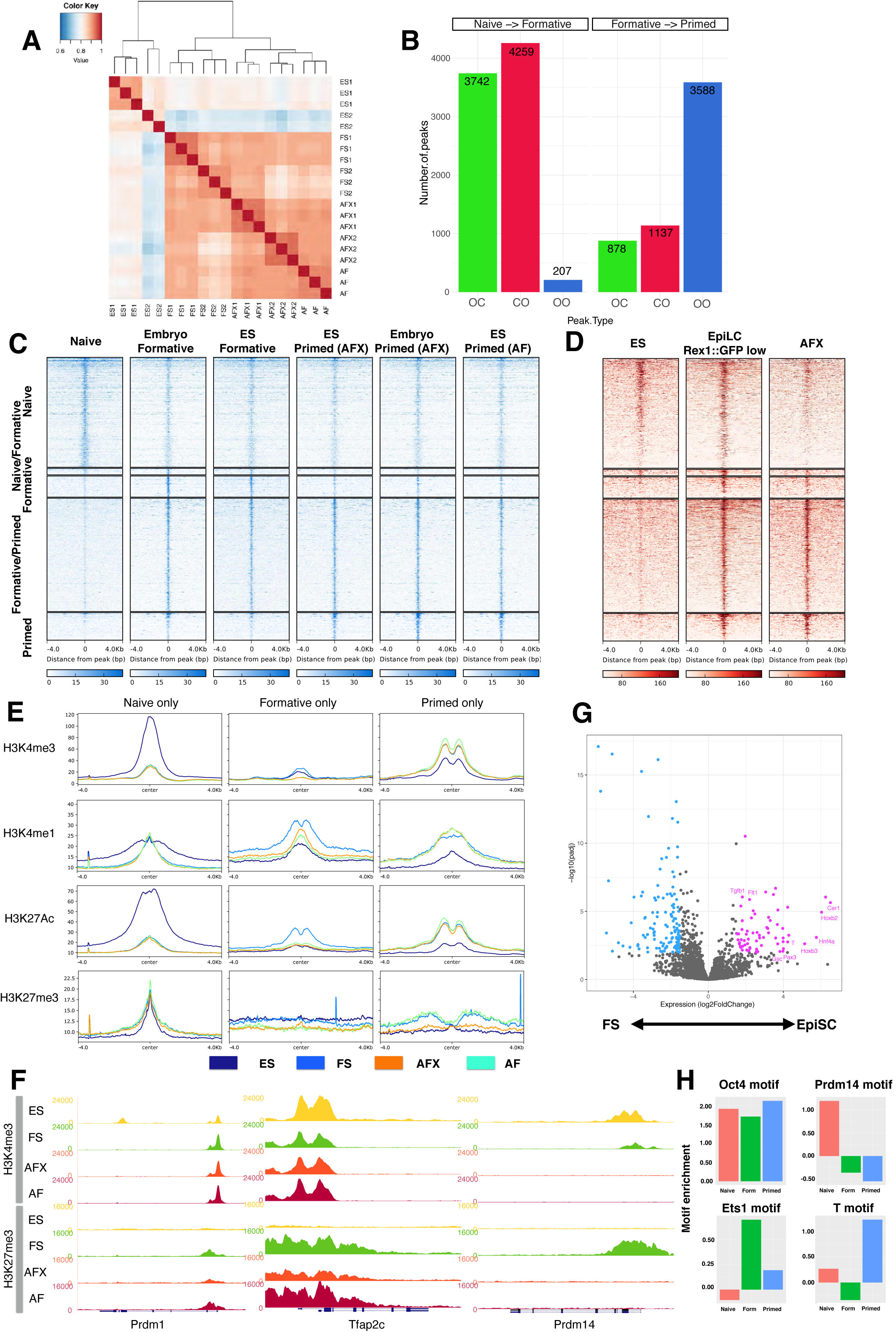
(A) Hierarchical clustering of all ATAC-seq peaks. (B) Number of specific peak changes between pluripotent states. OC; open to closed, CO; closed to open, OO; open to open. (C) Phase specific and shared ATAC-seq peaks. (D) Formative specific peaks identified in (C) are also enriched in transient EpiLCs. (E) Histone modification patterns at ATAC-seq peaks. (F) Genome browser screenshots of H3K4me3 and H3K27me3 distribution at *Prdm1, Tfap2c* and *Prdm14* loci. (G) Volcano plot shows gene expression fold changes between FS and EpiSCs from shared ATAC-seq peaks. Purple circles up-regulated in EpiSCs, blue circles up-regulated in FS cells. (H) Transcription factor binding motif enrichments at ATAC-seq peaks.

ChIP-seq for histone modifications confirmed the expected correlation between open chromatin and active marks, H3K4me3, H3K4me1, and H3K27Ac (Fig.5E). Regions that were more open in naïve and formative cells showed marked enrichment for H3K4me3 and H3K27ac that was lost in EpiSCs. Interestingly, active marks were also more highly represented in FS cells than in ES cells at loci that opened only in EpiSCs. We also examined bivalent promoter regions associated with both H3K4me3 and H3K27me3 (Azuara et al., 2006; Bernstein et al., 2006). We enumerated 2417 bivalent promoters in FS cells, nearly three times the number in ES cells (Fig. S5B). Many, but not all, of these loci were also bivalent in EpiSCs. Figure S5C shows examples of different profiles. Among the FS cell specific bivalent promoters was *Prdm14*, encoding one of the key germ cell determination factors (Nakaki et al., 2013). Promoters for other germ cell genes *Tfap2c* and *Prdm1* are also bivalent in FS cells, consistent with being poised for expression (Fig. 5F). In EpiSCs, however, *Prdm14* loses both marks indicating the gene is inactivated. This chromatin change may be related to the loss of competence for PGCLC induction in EpiSCs (Hayashi et al., 2011)

In addition we assessed DNA methylation at open chromatin regions using published data for EpiLCs and EpiSCs (Zylicz et al., 2015). In EpiLCs, all ATAC peaks were hypomethylated. In EpiSCs, however, naïve and formative peaks gained methylation, and only primed peaks maintained low methylation (Fig. S5D).

Interestingly, we observed marked differential expression between FS cells and EpiSCs among genes associated with shared ATAC peaks (Fig. 5G). Those more highly expressed in EpiSCs include gastrulation-associated factors such as Cer1, Gsc, T, Pax3. GO term analysis identifies enrichment for heart development, multicellular organism development and gastrulation (Fig. S5E). FS cell enriched transcripts are more numerous but comprise genes without annotated functions in early development (Table S1).

Using HOMER (Heinz et al., 2010) we identified transcription factor binding motifs enriched in open chromatin regions (Table S2). Core pluripotency factor binding motifs for Oct4 and Oct4-Sox-Tcf-Nanog are over-represented in all three cell types. ES cell ATAC peaks are also enriched for Tfcp2l1 and Prdm14 motifs, while those in EpiSCs feature Gsc, Brachyury, Slug, and Eomes motifs (Fig. 5H and S5F). Both FS cells and EpiSCs show increased accessibility of AP1/Jun sites. We noted that FS cell open chromatin shows specific enrichment for ETS-domain factor binding motifs.

### FS cells and EpiSCs show contrasting dependencies on Etv and Otx2

Previously we presented evidence linking Etv5, an ETS factor of the PEA3 sub-family, to enhancer activation during pluripotency progression (Kalkan et al., 2019b). We also showed that ES cells lacking Etv5 show diminished ability to make EpiSCs. Here we employed CRISPR/Cas9 to generate ES cells deficient for both Etv5 and the related Etv4. *Etv4/5*-dKO cells failed completely to produce EpiSCs upon transfer to AFX and differentiated into fibroblast-like cells (Fig. S6A). This phenotype is more severe than for *Etv5* mutation alone. Somewhat unexpectedly, however, *Etv4/5*-dKO cells converted to epithelial culture in A_lo_XR and subsequently expanded, albeit with persisting differentiation (Fig. 6A and S6A). Relative to ESCs, naïve factors are down-regulated and post-implantation markers up-regulated, including several direct targets of Etv5 such as Fgf5, Otx2 and Oct6 (Fig. 6B). We detected no compensatory up-regulation of the third PEA3 member, Etv1. *Etv4/5-*dKO FS cells differentiate readily via embryoid bodies and in directed protocols (Fig. S6B-E), including induction of Blimp1^+^, Stella^+^, Oct4^+^ PGCLC (Fig. S6F). However, when transferred to AFX, *Etv4/5-*dKO cells failed to convert to EpiSCs, lost expression of Oct4 within 3 days, and differentiated into fibroblasts with aberrant expression of Pou3f1 (Fig. 6C, D and Fig. S6G). Introduction of an *Etv5* transgene to *Etv4/5-*dKO cells restored the ability to convert to EpiSCs (Fig. 6E-H). These results establish that Etv4 and Etv5 are not essential for lineage competence of FS cells yet are required for production of EpiSCs *in vitro*.

**Fig. 6.**
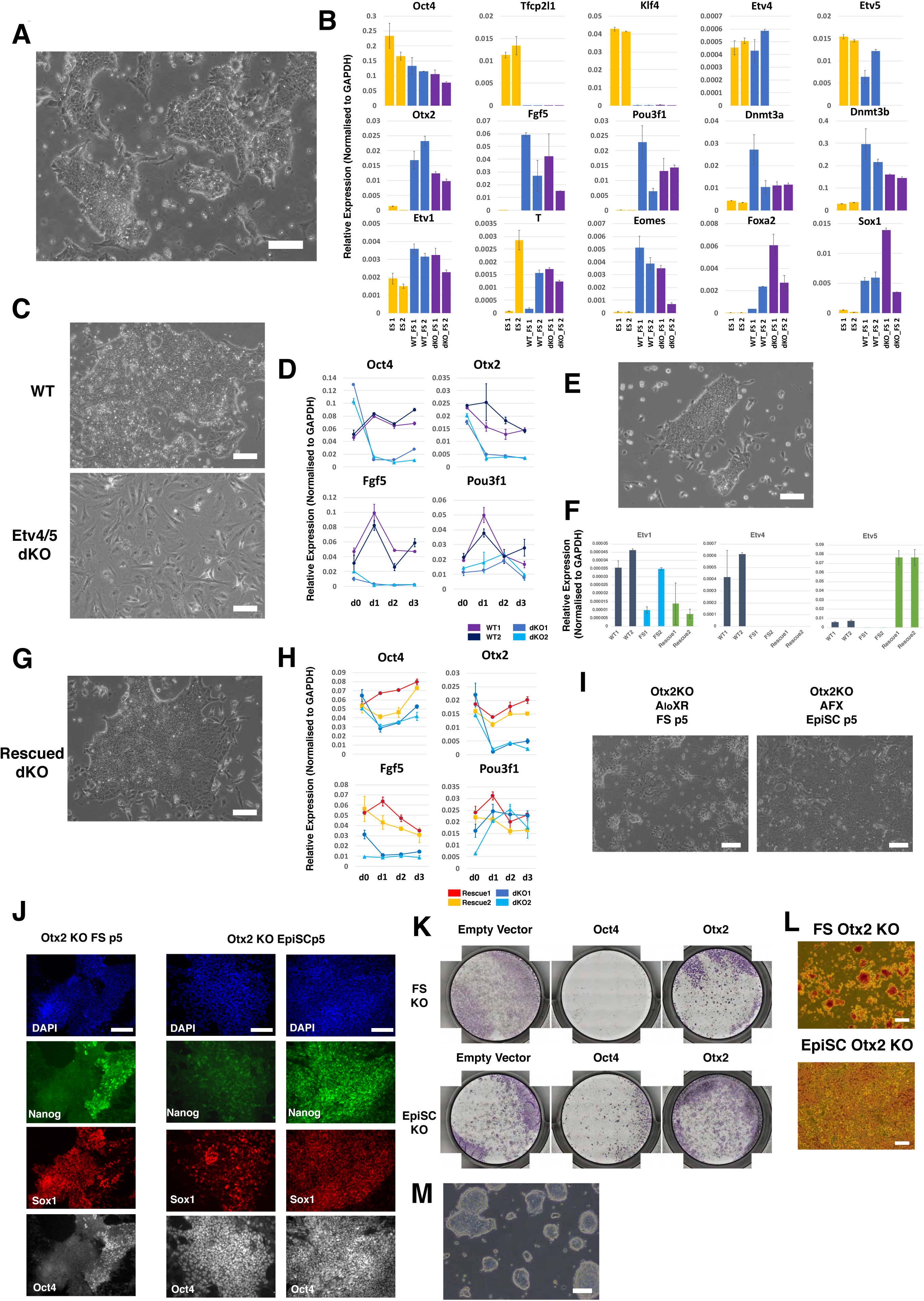
(A) Morphology of *Etv4/5* dKO FS cells. (B) RT-qPCR analysis of *Etv4/5* dKO FS cells. Two each of WT ES cells (yellow bars), WT FS cells (blue bars) and *Etv4/5* dKO FS cells (purple bars) were analysed. Error bars represents S.D. from technical duplicates. (C) Morphology of WT and dKO FS cells in EpiSC (AFX) culture medium for three days. (D) Time course RT-qPCR analysis of WT and *Etv4/5* dKO FS cells in EpiSC (AFX) culture. Error bars are S.D. from technical duplicates. (E) Morphology of *Etv4/5* dKO FS cells expressing *Etv5* transgene. (F) RT-qPCR profile of *Etv1*, -*4* and -*5* in *Etv5* rescue dKO lines. Error bars represents S.D. from technical duplicates. (G) Morphology of rescued dKO FS cells in EpiSC (AFX) culture. (H) Time course RT-qPCR analysis of rescued lines. Error bar represents S.D. from technical duplicates. (I) *Otx2* KO ES cells transferred to FS cell or EpiSC (AFX) culture conditions. Morphology at passage 5 shows massive neural differentiation observed in FS cell but not in EpiSC culture. (J) Immunostaining of *Otx2* KO cells at p5 in FS cells or EpiSC culture. Two classes of EpiSC colony were observed: left, homogenous Oct4 with heterogenous Nanog and Sox1; right, uniform Oct4, Sox1 and Nanog triple positive. (K) AP-staining of control, *Oct4* and *Otx2* KOs generated by Cas9/gRNA transfection in FS cells and EpiSCs. Colonies were stained three days after replating transfected cells. (L) Morphology of AP positive *Otx2* KO FS cells and EpiSCs. (M) Representative image of *Otx2* KO FS cells before the culture collapsed. Scale bars 100 µm, except (J) 50µm.

Otx2 is prominently up-regulated early during formative transition in vivo and in vitro (Acampora et al., 2016; Kalkan et al., 2017), and is implicated in redirecting genome occupancy of Oct4 (Buecker et al., 2014; Yang et al., 2014). Interestingly, *Otx2* is dispensable in both ES cells and EpiSCs (Acampora et al., 2013), but homozygous embryo mutants exhibit severe gastrulation phenotypes (Ang et al., 1996). We generated *Otx2* KO ES cells and investigated conversion into FS cells in A_lo_XR. Epithelial colonies emerged and could be expanded for 4-5 passages but continuously differentiated into neural cells (Fig. 6I). By passage 5 Oct4 and Nanog expression were downregulated and the majority of cells were positive for Sox1 (Fig. 6J). Cultures could not be maintained stably thereafter.In contrast Otx2 mutant ES cells could be converted into stable Oct4 positive EpiSCs in AFX (Fig. 6I), although colonies frequently displayed aberrant expression of Sox1 as previously reported (Acampora et al., 2013)(Fig. 6J). BMP has been shown to enhance stability of *Otx2* deficient EpiSCs (Acampora et al., 2013). We added BMP to two mutant FS cell cultures in A_lo_XR but observed no suppression of differentiation (Fig. S6H).

We also mutated *Otx2* directly in FS cells and observed that colonies became compact and dome-shaped, appearing rather similar to ground state ES cells (Fig. 6K, L, M). When replated in 2iL, however, *Otx2* mutant cells did not form ES cell-like colonies but differentiated or died (Fig.S6I). We repeated the targeting and achieved initial clonal expansion in A_lo_XR, but 8 out of 8 clones subsequently underwent massive neural differentiation and could not be stably propagated. We added BMP to three cultures, but this did not result in stabilisation. These results indicate that Otx2 is required for a stable FS cell state.

### Generation of human FS cells

We explored the possibility of deriving FS cells from naïve human pluripotent stem cells (hPSC) (Takashima et al., 2014). We used both chemically reset lines, cR-H9EOS and cR-Shef6 (Guo et al., 2017), and embryo-derived HNES cells (Guo et al., 2016). A_lo_X and A_lo_XR were applied as for mouse FS cell culture, except that plates were coated with a combination of laminin and fibronectin to improve attachment. The domed naïve hPSC converted to a more flattened epithelioid morphology over several days. Cultures could be propagated continuously thereafter and exhibited a faster doubling rate than naïve cells, requiring passage every 4 days at a split ratio of 1/15 (Fig. 7A). Cells in A_lo_XR exhibit loss of naïve markers (KLF4, KLF17, TFCP2L1) and retention of core pluripotency factor OCT4, but show little or no up-regulation of lineage priming markers, TBXT or FOXA2, often detected in conventional hPSC (Fig. 7B) (Allison et al., 2018; Gokhale et al., 2015). They show gain of SOX11 and OTX2, markers of post-implantation epiblast in the primate embryo (Nakamura et al., 2016).

**Fig. 7.**
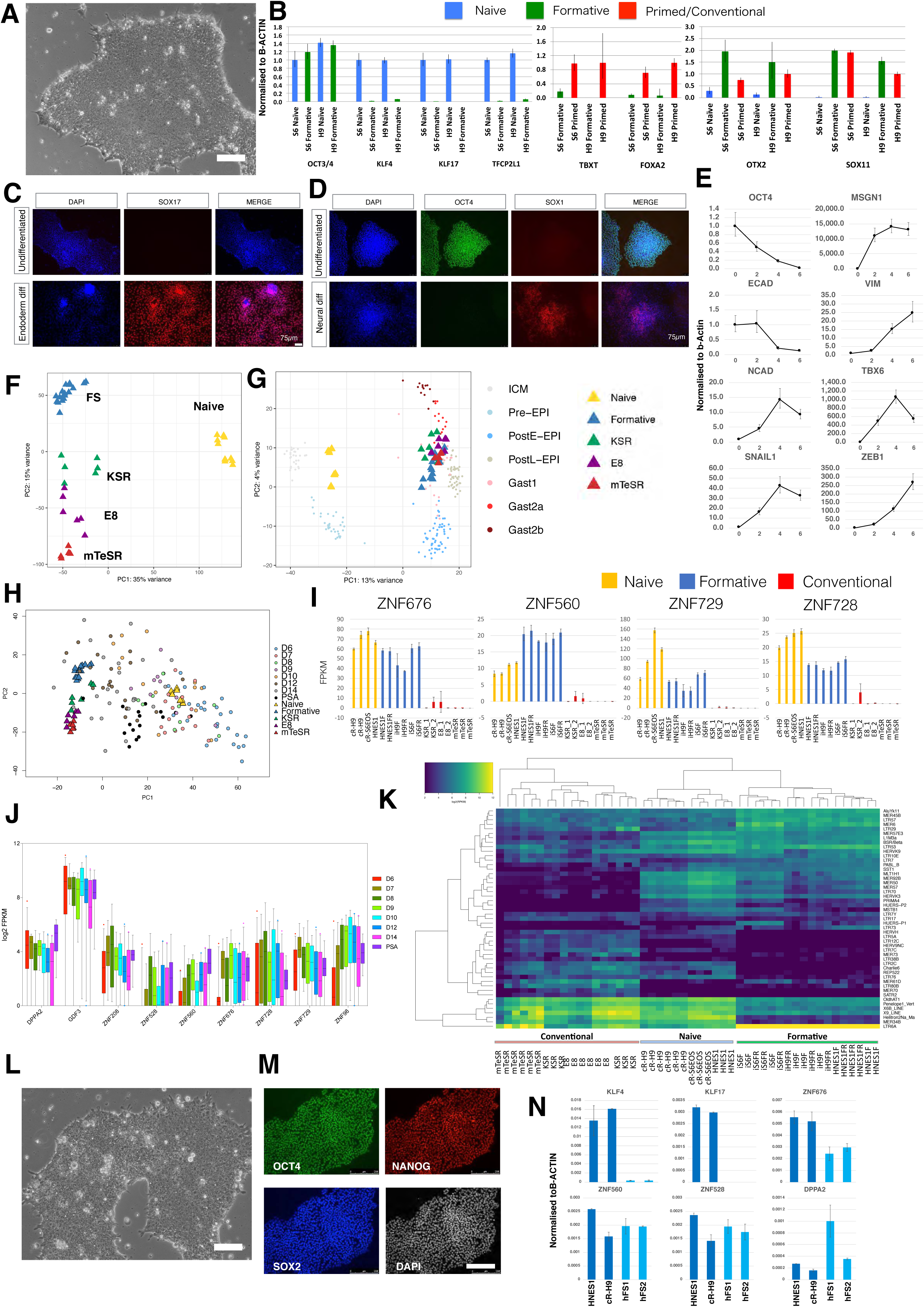
(A) Morphology of human FS cells derived from naïve hPSCs. Scale bar, 100µm. (B) Gene expression analysis of two hFS cell lines showing lack of naïve cell gene expression, low lineage marker gene expression and post-implantation related gene expression. Error bars represents S.D. from technical triplicates. (C) hFS cells differentiate into SOX17 positive endoderm cells. (D) hFS cells differentiate into SOX1 positive neuroepithelial cells. (E) RT-qPCR analysis of hFS cells differentiated into paraxial mesoderm cells. Error bars represent S.D. from technical triplicates. (F) PCA of hFS cells with naïve and conventional hPSCs computed with 11051 genes identified by median Log2 expression >0.5. (G) Projection of human FS cell and conventional PSC samples onto PCA of *Macaca* ICM/epiblast stages computed with 9432 orthologous expressed genes. (H) PCA for cell line populations with human embryo single cells projected. PCA computed using 922 variable genes across epiblast samples from human embryo extended culture (Xiang et al., 2019). (I) FPKM values for naïve-formative specific genes in naïve, formative or conventional hPSCs. (J) Boxplots of naïve-formative specific gene expression in human epiblast stages and PSA (K) Heatmap analysis of differentially expressed transposable element analysis between naïve, formative and conventional samples. (L) Morphology of FS cells derived directly from human embryo. Scale bar, 100µm. (M) Immunostaining of OCT4, SOX2 and NANOG in embryo-derived hFS cells. Scale bar, 250µm (N) RT-qPCR analysis of embryo-derived hFS cells. Error bars represent S.D. from technical duplicates.

Naïve hPSC do not respond productively to somatic lineage induction protocols but must first undergo formative transition to lineage competence (Guo et al., 2017). This capacitation process takes place over several days (Rostovskaya et al., 2019). FS cells in contrast are expected to be directly responsive to lineage cues. We applied established protocols for differentiation to human FS cells. In response to definitive endoderm induction (Loh et al., 2014), we observed efficient formation of SOX17 positive cells (Fig. 7C), while neural induction via dual SMAD inhibition (Chambers et al., 2009) resulted in abundant SOX1 immunopositive cells (Fig. 7D). We also tested paraxial mesoderm differentiation (Chal et al., 2016) and detected up-regulation of *TBX6* and *MSGN1* along with EMT markers such as *SNAIL1* and *ZEB1* (Fig. 7E).

We prepared RNA-seq libraries from three human FS cell lines and carried out whole transcriptome comparison with naïve and conventional hPSCs (Fig 7F). PCA separates naive cells on PC1 and distinguishes formative from conventional hPSC on PC2, similar to the analysis of mouse PSCs (Fig. 4A). Transcriptome data are not available for early post-implantation embryogenesis in human. We therefore took advantage of expression data for the non-human primate *Macaca fascicularis* (Nakamura et al., 2016). We computed the PCA for *Macaca* using 9324 expressed orthologous genes (median Log2 expression>0.5) onto which we projected the human cell line samples (Fig. 7G). FS cells and conventional hPSCs aligned with post-implantation embryo stages. FS cell samples aligned with late post-implantation epiblast while conventional hPSCs spread from late epiblast to early gastrulating cells.

Single cell transcriptome data has recently been published for human embryos during extended culture (Xiang et al., 2019). We used variable genes in the epiblast and primitive streak anlage (PSA) stages to compute the PCA for naïve, formative and conventional hPSCs onto which the embryo single cells were projected. The resulting plot shows a similar pattern to the *Macaca* embryo comparison; naïve cells clustered with pre-implantation epiblast and formative cells next to post-implantation stages with conventional hPSCs adjacent but distributed more towards the PSA cluster (Fig. 7H).

We performed K-means clustering between formative and conventional PSC cultures (k=6) (Fig S7A). Cluster 1 comprises 369 genes expressed more highly in FS cells than conventional hPSCs. The majority of protein-coding genes in this cluster are expressed in naïve cells and persist during capacitation (Fig S7B, C). *DPPA2, GDF3* and several *ZNF* genes were identified as useful markers expressed in both naïve and formative cells but variably low or absent in conventional hPSCs (Fig. 7I, S7D). Expression of these ZNF genes was detected in human pre- and post-implantation epiblast transcriptome data (Fig. 7J).

KRAB-ZNFs such as ZNF676, ZNF560, and ZNF528 can suppress expression of transposable elements (TEs) (Friedli and Trono, 2015). TEs are dynamically expressed in early development and highly differential between naïve and primed hPSC (Friedli and Trono, 2015; Guo et al., 2017; Theunissen et al., 2016). We examined TE expression in FS cells and observed a distinct profile compared with naïve or conventional hPSCs (Fig. 7K). For example, FS cells distinctively express LTR6A, and retain expression of certain HERVK TEs also expressed in naïve cells, but do not express subsets of SVAs family members that are prominent in naive cells, or subsets of HERVH, LTR7C or LTR12C family members that are prominent in primed cells (Fig. S7E).

Finally, we investigated application of FS cell culture conditions directly to human ICMs. We thawed E5 and E6 embryos and cultured for one or two days respectively in N2B27. We then isolated ICMs by immunosurgery or mechanical dissociation and plated them intact on laminin/fibronectin coated dishes in A_lo_XR with ROCK inhibitor Y-27632. After 2-4 weeks, primary outgrowths were manually dissociated and re-plated. We established three lines from different embryos. The embryo derived lines exhibited similar morphology and growth behaviour to naïve PSC derived FS cells (Fig. 7L). G-banded karyotype analysis showed that all three expanded lines are diploid (46XX, 20/20) (Fig.S7G). We confirmed relatively homogeneous expression of OCT4, SOX2 and NANOG by immunostaining (Fig. 7M). Expression of naïve-specific transcription factors KLF4 and KLF17 was not detected while transcripts were present for several genes that are expressed in naïve and formative cells but down-regulated in conventional hPSCs (Fig 7N).

## DISCUSSION

Expandable stem cells that retain high fidelity to staging posts of pluripotency in the embryo will be instrumental for harnessing capacity to recapitulate development, create disease models, and manufacture therapeutic cells. Stem cells representative of naïve and primed pluripotency have been established in mouse and human (Davidson et al., 2015; Nichols and Smith, 2009; Rossant, 2015; Rossant and Tam, 2017) but formative pluripotency has only been characterised in the form of transient EpiLCs (Buecker et al., 2014; Hayashi et al., 2011; Kalkan et al., 2017; Mulas et al., 2017). In this study we attempted to fill the stem cell void between early and late pluripotency (Kinoshita and Smith, 2018; Smith, 2017). We show that in defined culture conditions inhibition of Wnt and retinoid signalling in combination with low levels of activin enables capture of stem cells that retain key features of formative stage mouse epiblast. FS cells have exited the ES cell state and diverged comprehensively in transcription factor repertoire, chromatin profile, and culture requirements. They are maintained by FGF and nodal/activin but are poised to respond to increased levels of either signal by initiating mesendoderm differentiation. FS cells are distinguished functionally from EpiSCs by the abilities to generate germline precursors and to colonise chimaeras following blastocyst injection. At the transcriptome level FS cells are related to pre-streak formative epiblast and are largely devoid of lineage marker expression. They show specific transcription factor dependencies from either ES cells or EpiSCs. Finally, FS cell culture conditions applied to human naïve PSCs or embryo explants support expansion of stem cells related to post-implantation epiblast without use of feeders, KSR or FGF, standard components of conventional hPSC culture.

Previously various culture conditions have been reported to support propagation of mouse ES cell derivatives with features of late blastocyst or peri-implantation epiblast, such as reduced expression of Rex1 or increased expression of Otx2 (D’Aniello et al., 2016; Neagu et al., 2020; Rathjen et al., 1999). However, these cells spontaneously revert to the canonical ES cell phenotype when transferred back to ES cell culture. Therefore, they remain within the ES cell spectrum. Similarly, reversible down-regulation of naïve pluripotency factors and partial differentiation is observed in ES cell cultures in serum (Chambers et al., 2007; Filipczyk et al., 2015; Hayashi et al., 2008; Toyooka et al., 2008). The cytokine LIF, which potently promotes mouse ES cell identity (Dunn et al., 2014; Smith et al., 1988; Williams et al., 1988), is a key component of all these culture conditions. In contrast, FS cells are maintained without LIF and show no reversion to ES cell phenotype. Therefore, FS cells have extinguished naïve epiblast character, in line with loss of ability for ES cell formation in early post-implantation epiblast (Boroviak et al., 2014).

In mouse, a defining functional attribute of formative epiblast is direct responsiveness to germline induction, which is lacking in both naïve cells and primed gastrula stage epiblast (Ohinata et al., 2009). Differentiation of ESCs into transient EpiLC populations recapitulates a brief window of germline competence but this last only from 24-72 hrs (Hayashi et al., 2011). Maintenance of germline competence together with somatic competence over many passages is therefore a unique feature of mouse FS cells.

Mouse FS cells also differ from ES cells and EpiSCs in their contribution to chimaeras. Chimaerism is less frequent, to lower levels, and less evenly distributed than typically obtained with ES cells. Poorer contributions are not unexpected given the heterochronicity between FS cells and E3.5 host blastocysts. Indeed, the pioneering mouse embryo chimaera studies suggested that blastocyst colonisation capacity was completely lost after implantation (Gardner, 1985). Here, using more sensitive detection systems we show that after formation of the pro-amniotic cavity epithelialised formative epiblast cells can still contribute to blastocyst chimaeras, in a comparable manner to FS cells. EpiSCs in contrast do not generally show any significant contribution to chimaeras via blastocyst injection, unless they have been genetically engineered (Masaki et al., 2016; Ohtsuka et al., 2012; Tesar et al., 2007). Intriguingly, it has been reported that certain EpiSC lines cultured on feeders or serum-coated dishes contain a sub-population of cells that are able to contribute to chimaeras (Han et al., 2010; Kurek et al., 2015). The nature of such cells is unclear, but our results raise the possibility that they may represent FS cells co-existing with EpiSCs in those undefined conditions.

FS cells exhibit distinct signal dependency and responsiveness compared to ESCs or EpiSCs. Both mouse EpiSCs and human conventional PSCs are cultured in medium supplemented with FGF. Indeed, high FGF (100ng/ml) is considered an essential component of defined E8 medium for hPSCs (Chen et al., 2011; Cornacchia et al., 2019). FS cells in contrast are cultured without addition of FGF. Notably mouse FS cells respond directly to FGF and other stimuli for primitive streak induction by up-regulating *T*. Consistent with readiness for *T* induction, FS cells exhibit greater propensity to form mesendoderm than EpiSCs. Neural lineage entry, on the other hand, is delayed by around 24hrs, but the final differentiation efficiency appears similar. These observations are consistent with FS cells representing pre-streak epiblast with relatively unbiased responsiveness to primitive streak and neural induction. We surmise that the relative recalcitrance of EpiSCs to primitive streak induction may reflect adaptation to the high growth factor signals that drive their in vitro proliferation.

Whole transcriptome analysis substantiates that mouse FS cells are distinct from EpiSCs but related to EpiLCs. Compared with the embryo they are most similar to E5.5-6.5 egg cylinder epiblast, the stage of formative pluripotency. Single cell analysis shows that FS cells are more uniform than EpiSCs. In human, FS cells appear related to a somewhat earlier stage of post-implantation epiblast than conventional hPSCs. However, conventional hPSCs are not equivalent to EpiSCs (Lau et al., 2020), and the differences from FS cells are less marked.

Open chromatin regions differ between ES and FS cells, consistent with reconfiguration to generate a chromatin platform poised for lineage specification. By contrast, conversion of FS cells to EpiSCs is accompanied by fewer chromatin changes, in line with continuous development. Nonetheless we identified distinct transcription factor dependencies between FS cells and EpiSCs. FS cells are only mildly destabilised by deletion of Etv5 and Etv4 and remain pluripotent, whereas the EpiSC state cannot be established in the absence of these factors (Kalkan et al., 2019b). Whether the inability to produce *Etv4/5* dKO EpiSCs reflects a specific functional requirement in EpiSCs or results from a cryptic change in formative competence remains to clarified. Interestingly, at least a proportion of *Etv5* or *Etv4/5* mutants proceed through gastrulation to E8.5 or E9.5 (Lu et al., 2009; Zhang et al., 2009). The *Etv4/5* knockout phenotypes suggest that the *in vitro* EpiSC state may not correspond closely to a waystation of epiblast progression *in vivo* (Kojima et al., 2014). Conversely, we found that Otx2 is indispensable for stable expansion of FS cells. Otx2 is not required by ES cells or EpiSCs (Acampora et al., 2013) but is essential for *in vivo* gastrulation (Ang et al., 1996). EpiSCs lacking Otx2 do exhibit aberrant features (Acampora et al., 2013), however, which we speculate may be a consequence of defective formative transition.

In FS cells the transcription factor circuitry governing naïve pluripotency (Dunn et al., 2014; Takashima et al., 2014) is dismantled, signalling pathways are rewired, and chromatin accessibility is extensively remodelled, consistent with a discontinuous change in competence as cells differentiate from naïve to formative states. By contrast, the molecular separation between FS cells and primed pluripotent stem cells is less distinct in line with more continuous developmental progression. We surmise that the reconstructed gene regulatory network and chromatin landscape in formative cells provides the requisite context for signalling cues to induce germ layer and germline lineage specification and for the subsequent unfolding of gastrulation. Capture of formative phase cells as self-renewing stem cell cultures should be enabling for comprehensive interrogation of the molecular features that confer and effect multi-lineage potency.

### Limitations of Study

Although the formative phenotype is produced within 48hrs of ESC withdrawal from 2i, generation of stable FS cell lines requires several passages. The inherent asynchronicity of exit from naïve pluripotency (Strawbridge et al., 2020) together with imperfect in vitro transition conditions result in extensive initial heterogeneity, as also observed for EpiLC (Hayashi et al., 2011; Kalkan et al., 2017). Passaging enriches for FS cells, similar to EpiSC generation (Guo et al., 2009), but a more streamlined and efficient capture would be advantageous for future research. In mouse, FS cells are clearly distinguished from EpiSCs by several features, most notably competence for germ cell induction and ability to colonise chimaeras via blastocyst injection. Neither of those functional criteria are applicable in the human context. Conventional hPSCs share some features with EpiSCs but do not appear to be direct equivalents (Lau et al., 2020; Rossant and Tam, 2017). Notably they can be induced to form primordial germ cell-like cells (Irie et al., 2015; Sasaki et al., 2015). Chimaera contribution cannot be tested in human embryos. At the transcriptome level human FS cells differ from populations of conventional hPSCs cultured in E8 or other conditions, but these differences are relative rather than absolute. Heterogeneity and hierarchical substructure has been described in hPSC cultures (Allison et al., 2018; Hough et al., 2009; Hough et al., 2014; Lau et al., 2020; Nakanishi et al., 2019) and we cannot exclude the presence of formative stem cells at some frequency. Human FS cells and conventional hPSCs may be a continuum spanning stages of post-implantation epiblast. It will be valuable in future studies to define marker sets and in vitro differentiation features that can better distinguish human formative cells from downstream stages in the spectrum of post-naïve pluripotency. To this end additional transcriptomic and other data from post-implantation epiblast will be important to allow more precise comparison and staging.

## Acknowledgements

We thank Vicki Metzis for the ATAC-seq protocol, Elsa Sousa for help with *Xist* FISH, and Meng Amy Li for the gRNA expression vector. Tüzer Kalkan, Nicola Reynolds and Laurence Bates advised on ChIP-seq protocols. We are grateful to Maike Paramor, Vicki Murray, Peter Humphreys, Darran Clements, Andrew Riddell, Charles-Etienne Dumeau and biofacility staff for technical support and to the CSCI core bioinformatics team for data processing and routine analysis. Sequencing was performed by the CRUK Cambridge Institute Genomics Core Facility. Shahzaib Ahmed and Benjamin Porteous contributed to experiments and Rosalind Drummond and James Clarke provided laboratory assistance. We thank Brian Hendrich for comments on the manuscript. This research was funded by the Biotechnology and Biological Sciences Research Council and the Medical Research Council of the United Kingdom. The Cambridge Stem Cell Institute receives core funding from Wellcome and the Medical Research Council. AS is a Medical Research Council Professor.

## Author contributions

Conceptualization, AS; methods, MK; formal analysis, MB, DS, GGS, SD; investigation, MK, WM, YC, JN; writing, MK, AS; supervision, AS

## Supplemental Figure Legends

**Figure S1, related to Figure 1.**
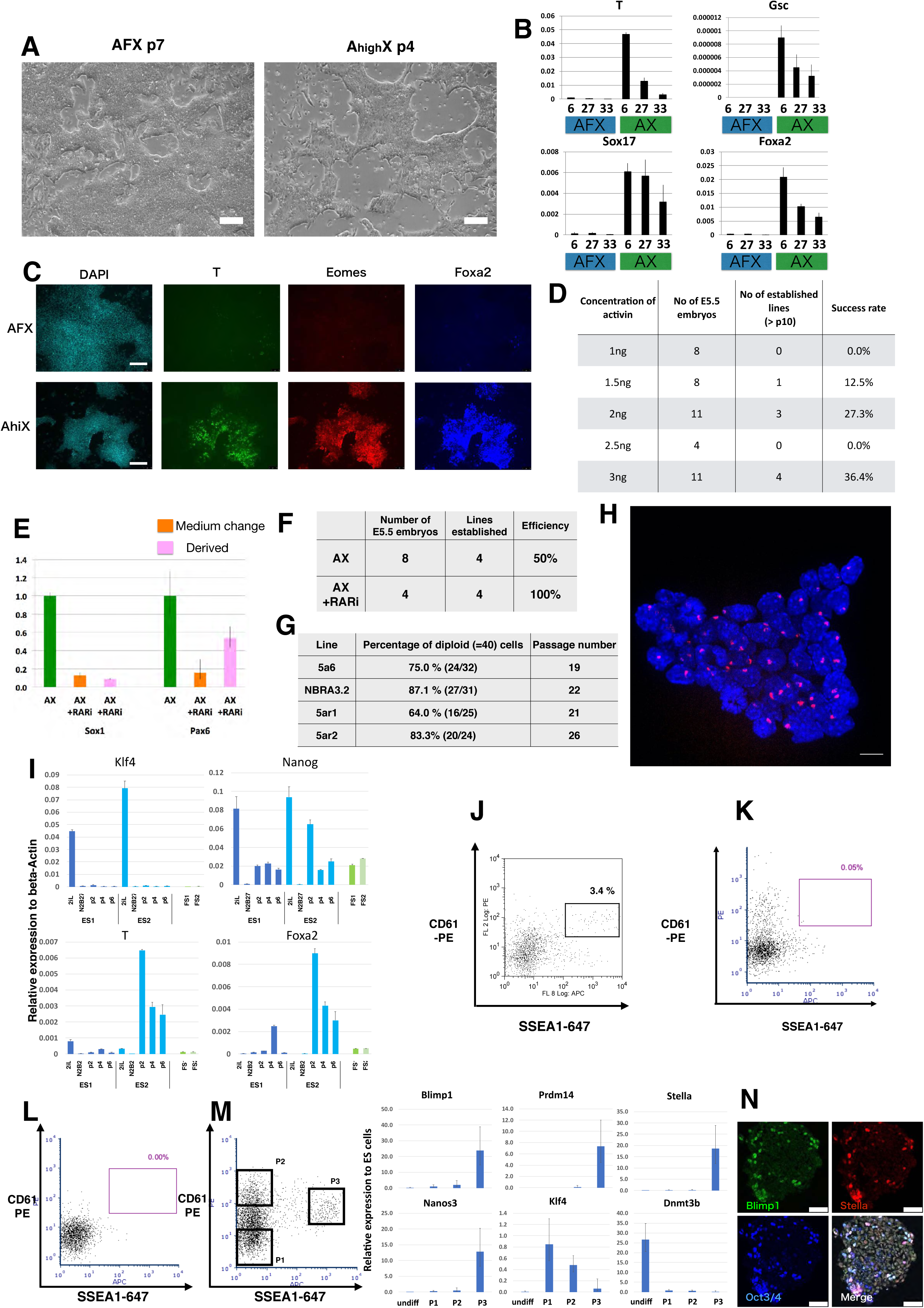
(A) Bright field image of E5.5 epiblast derived AFX colonies and A_hi_X colonies. Scale bars, 200µm. (B) Gene expression analysis of FGF withdrawal. Three AFX cell lines (6, 27 and 33) were passaged without FGF and analysed by RT-qPCR. Error bar represents S.D. from technical triplicate. (C) Immunostained images of AFX and A_hi_X cells showing lineage markers in A_hi_X cells. Scale bars, 100 µm. (D) Summary of derivation efficiency from E5.5 epiblasts in different concentrations of activin A. (E) RT-qPCR analysis of RAR inhibitor treated cells. A_lo_XR sample established in A_lo_X and cultured in A_lo_XR are shown in orange and a new line established from an E5.5 embryo in A_lo_XR in pink. Error bars, technical triplicates. (F) Derivation efficiency in the presence of RAR inhibitor from E5.5 epiblasts. (G) Percentages of diploid cells for 4 different FS cell lines. (H) Maximum projection of Z-stack slices of *Xist* FISH images (red) in female FS cells. Nuclei were stained with DAPI (blue). Scale bar, 10µm. (I) Gene expression analysis during ES cell to FS cell conversion. Gene expression is shown relative to beta-actin. Error bars are S.D. from two technical replicates. (J) Flow cytometry analysis of day 4 PGCLC induction from A_lo_X FS cells. (K) Analysis of day 4 PGCLC induction from AFX EpiSCs. (L) Analysis of day4 PGCLC induction from AFX EpiSCs adapted to culture in A_lo_XR. (M) RT-qPCR analysis of day 6 PGCLC from A_lo_XR cells sorted for SSEA1 and CD61 co-expression. Relative expression level to 2iL ES cells (=1) was normalized to Tbp. Error bars represent S.D. from technical triplicates. (N) Immunostaining of A_lo_X PGCLC. Scale bars, 50µm.

**Figure S2, related to Figure 2.**
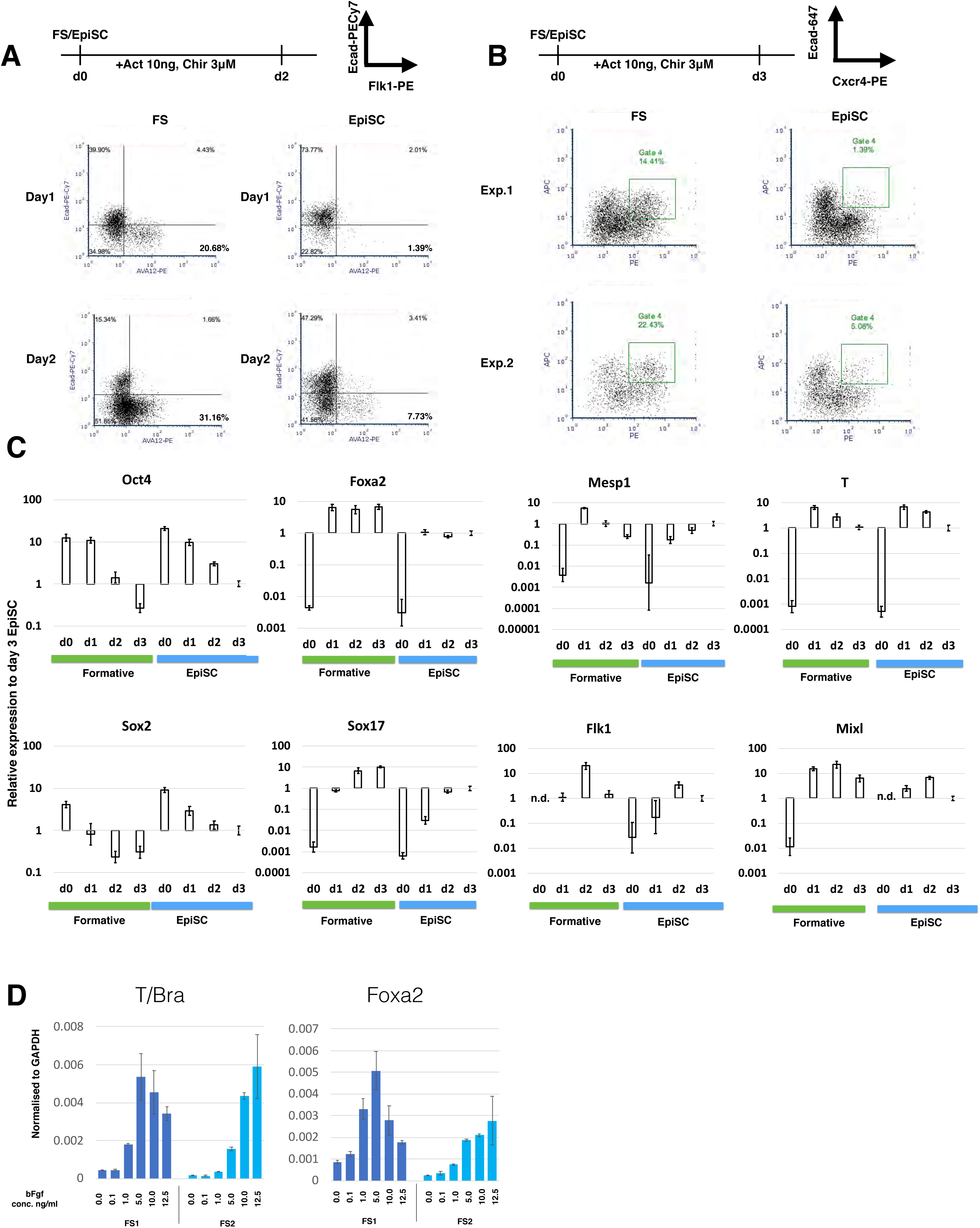
(A) Flow cytometry profiles of Flk1^+^Ecad^-^ mesodermal fraction of differentiated FS cells and EpiSCs at day 1 and day 2. (B) Cxcr4^+^Ecad^+^ endoderm fraction at day 3. Two experiments are shown. (C) RT-qPCR analysis after activin A and CHIR treatment for 3 days. AFX samples at day 3 were set as 1, normalised to 36B4 (Rplp0). Error bars represent SD from technical triplicates. n.d. indicated not detected. (D) RT-qPCR analysis of T and Foxa2 expression 24 hours after indicated doses of Fgf2 were added into A_lo_XR culture. Error bars represent S.D. from technical duplicates.

**Figure S3, related to Figure 3.**
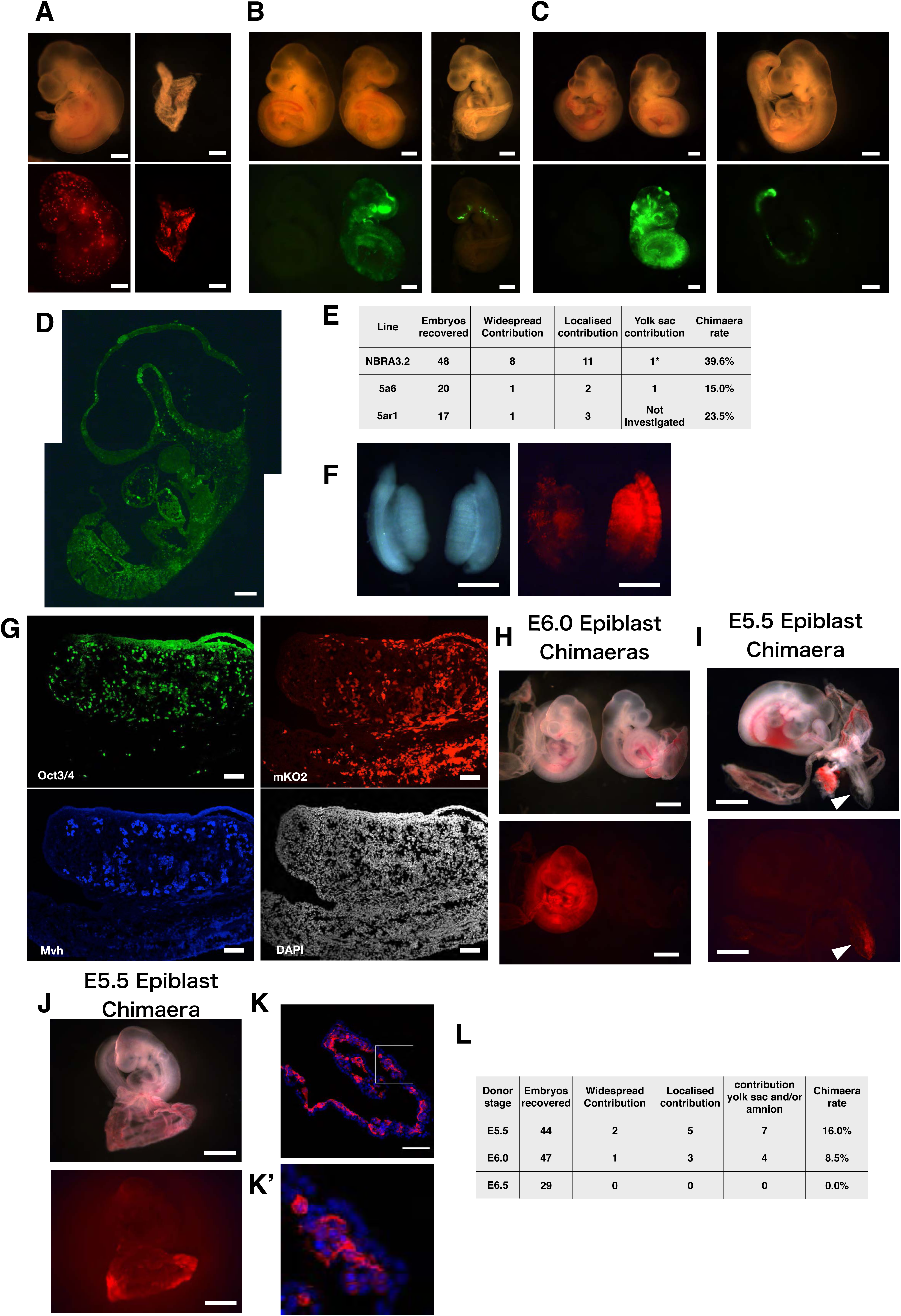
(A) Low contribution chimaera at E9.5 chimaera (left) produced from mKO2-labelled NBRA3.2 FS cells. mKO2 positive cell were found in yolk sac from one of the chimaeras shown in Fig. 3A (right). Scale bars, 500µm. (B) E9.5 chimaeras from GFP-labelled 5a6 FS cell line. Contributions were widespread (left) or localised (right). Scale bars, 500µm. (C) E9.5 chimaeras from GFP-labeled 5ar1 FS cells. Scale bars, 500 µm. (D) Sagittal section of embryo from C, left panel, showed widespread contribution of GFP positive cells. Scale bar, 200 µm. (E) Summary of FS cell chimaeras examined at E9.5. *Not all yolk sacs from chimaeric embryos were examined. (F) E12.5 chimaeric gonads generated from mKO2-labelled FS cells. Scale bars, 500µm. (G) Section of gonad from (F). Section was stained with anti-Oct4 and anti-Mvh antibodies. Nuclei were stained with DAPI. (H-J). E9.5 chimaeras with contribution from E5.5 and E6.0 donor epiblast. Contributions were detected in the embryo proper and yolk sac (H), amnion (arrowhead) (I), yolk sac (J). Scale bars, 1mm. (K) Yolk sac section. Membrane-tdTomato positive cells are present in the inner layer extraembryonic mesoderm. Nuclei were stained with DAPI (blue). Scale bar, 100µm. Magnified image from boxed region is shown as (K’). (L) Summary of post-implantation epiblast chimaeras.

**Figure S4, related to Figure 4.**
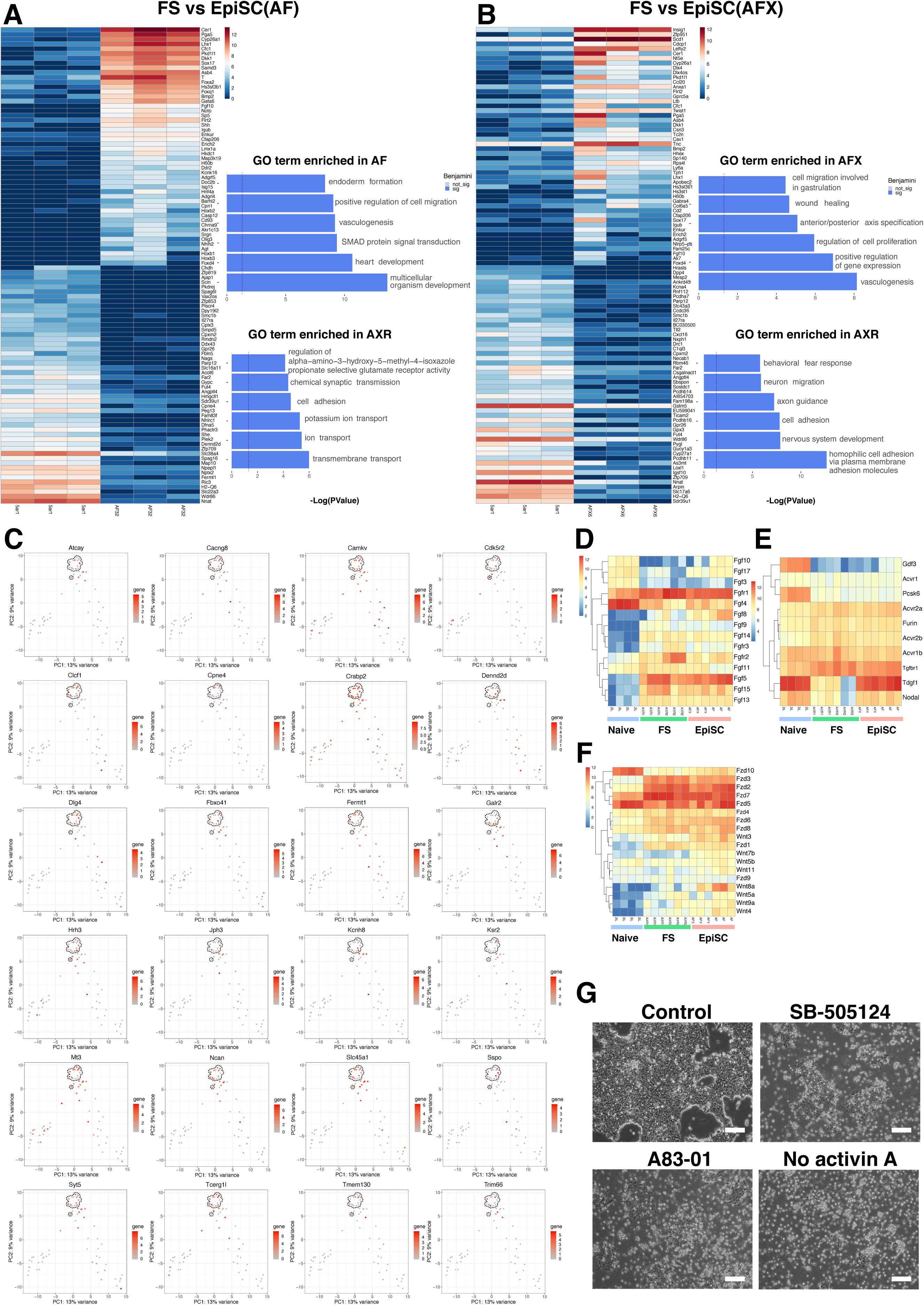
(A) Top 50 differentially expressed gene analysis between FS cells and EpiSCs (AF). GO term analysis was performed on 200 DEG. Top 6 GO terms are shown (benjamini value<0.05). (B) Top 50 differentially expressed gene analysis between FS cells and EpiSC (AFX) cells. GO term analysis as in A (benjamini value<0.05). (C) Examples of the gene expression pattern of FS cell specific genes identified in Fig. 4B. (D) Heatmap of expression of Fgfs and Fgfrs. (E) Heatmap of Nodal pathway gene expression. (F) Heatmap of expression of Wnts and Fzd receptors. Colour scale in (D-E) is log2(normalised counts +1) from RNA-seq. (G) Cell morphologies after two days in indicated culture conditions: A_lo_XR; A_lo_XR plus 1µM A83-01; A_lo_XR plus 5µM SB505124; XR without activin A. Scale bars, 100 µm.

**Figure S5, related to Figure 5.**
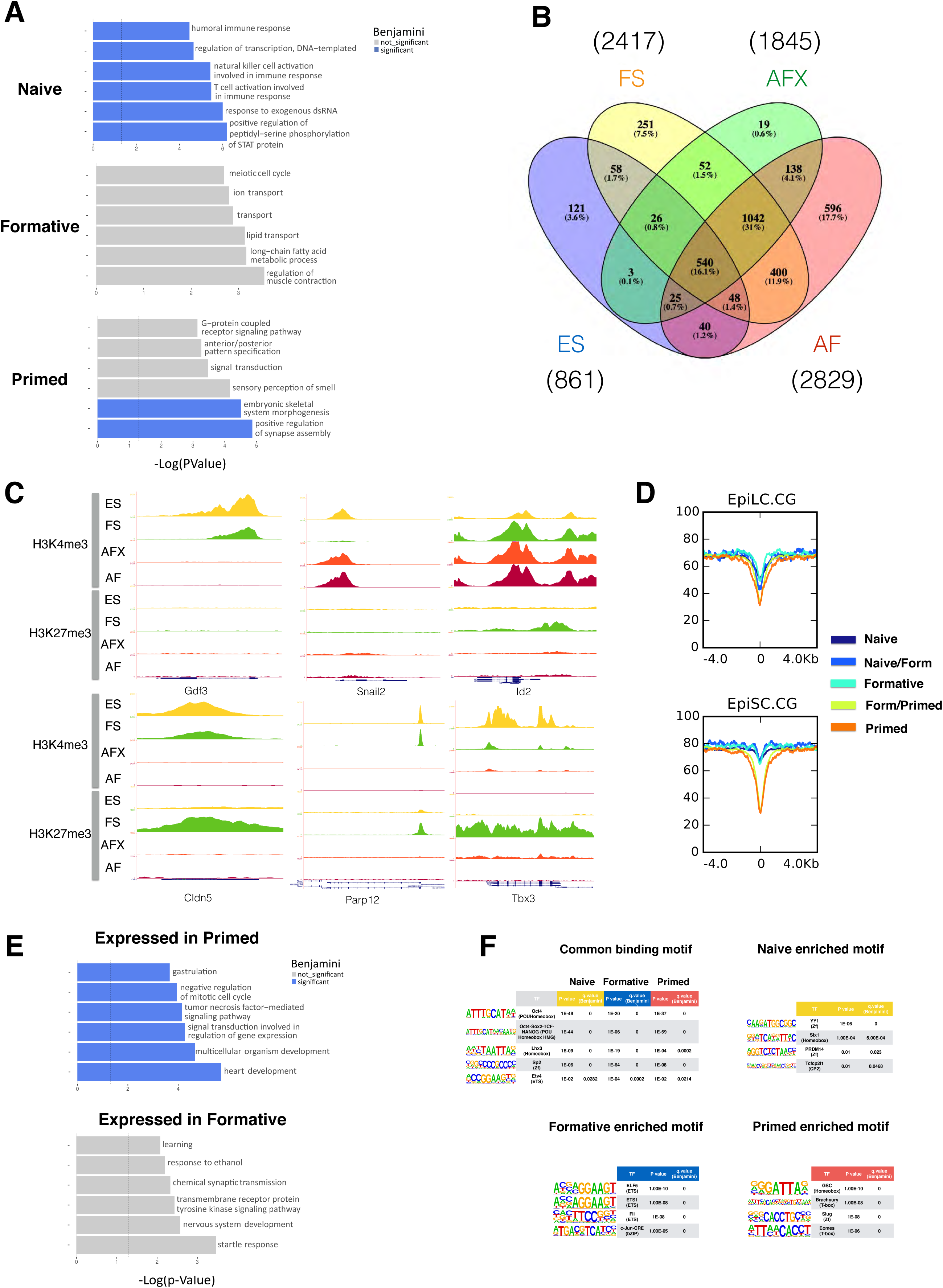
(A) GO term enrichment of phase specific ATAC-seq sites. Bars in blue have a significant benjamini value<0.05. (B) Venn diagram showing numbers of shared and unique bivalent domains in each cell type. (C) Genome browser examples of differential histone modifications. Lower three show formative specific bivalency. (D) Methylation at ATAC peaks in EpiLCs and EpiSCs (original data from Zylicz et al., 2015). (E) Related to Fig. 5G. GO term analysis for differentially expressed genes. Bars in blue have a significant benjamini value<0.05. (F) TF motif and P-values enriched in phase specific sites.

**Figure S6, related to Figure 6.**
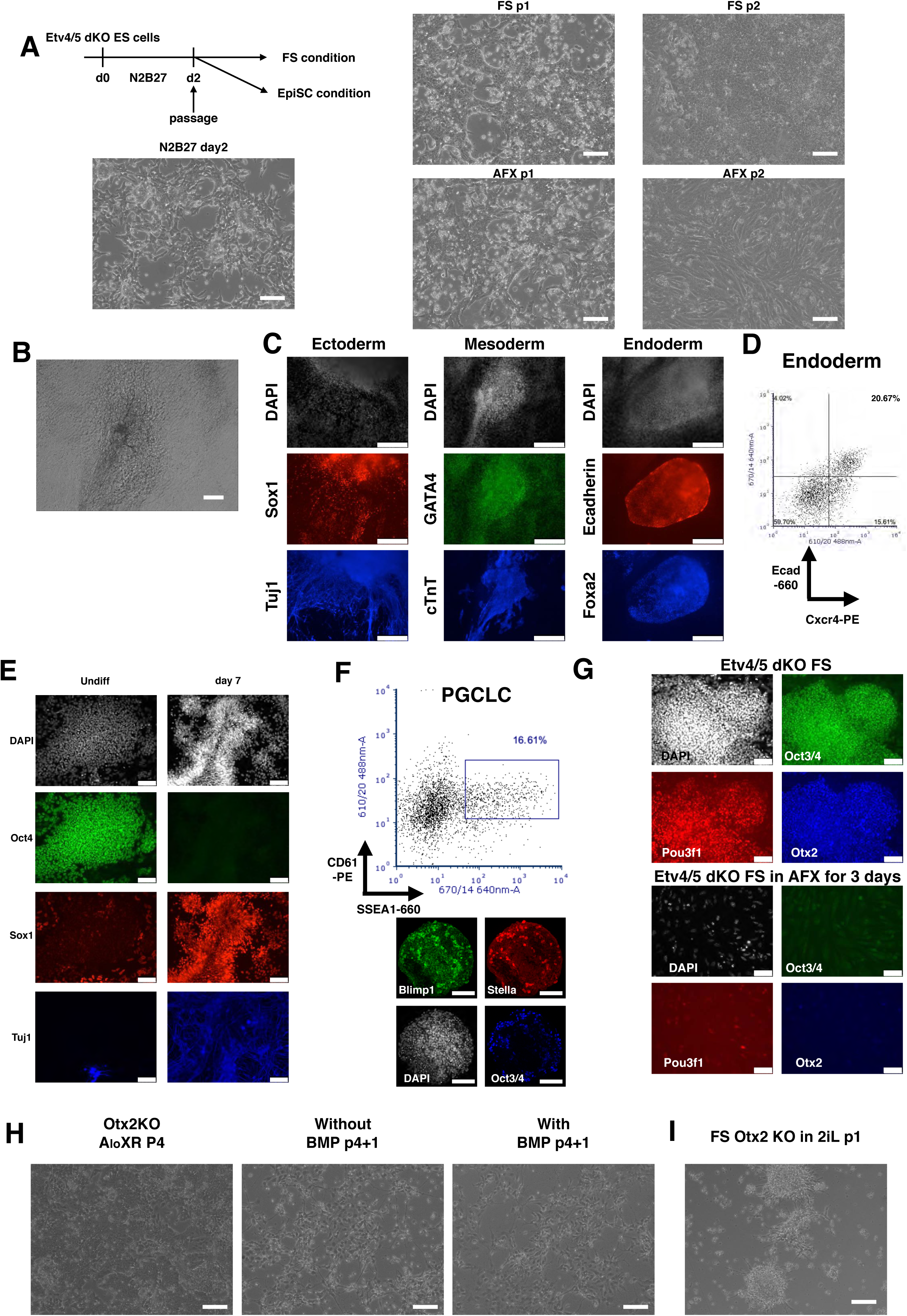
(A) Schematic of ES cell differentiation to FS cells or EpiSCs. Morphology at day 2, p1 and p2 are shown. (B) Bright field image of beating dKO derived cells. (C) Immunostaining of dKO FS cell EB outgrowth. Ectoderm cells were stained with Sox1 (red) and Tuj1 (Blue), Mesoderm cells were stained with Gata4 (Green) and cTnT (blue) and endoderm cells were stained with Ecadherin (red) and Foxa2 (Blue). DAPI stainings were shown in white. (D) FACS plot of endoderm differentiated dKO FS cells. Cxcr4-PE and Ecad-660 were used. (E) Neural differentiation of dKO FS cells. Immunostaining for Oct3/4 (green), Sox1 (red) and Tuj1 (Blue). (F) dKO FS cells subjected to PGCLC induction. Cells were analysed by flow cytometry for SSEA1-660 and CD61-PE and by immunostaining for Blimp1 (green), Stella (red) and Oct3/4 (blue). (G) Immunostaining of Etv4/5-dKO FS cells and after transfer to EpiSC culture for three days. (H) Otx2 KO cells passaged in A_lo_XR with or without BMP. (I) Bright field image of Otx2 KO FS cells re-plated in 2iL. Scale bars in (A), (B), (F), (H), (I) 100µm, (C) 250µm and (E), (G) 75µm.

**Figure S7, related to Figure 7.**
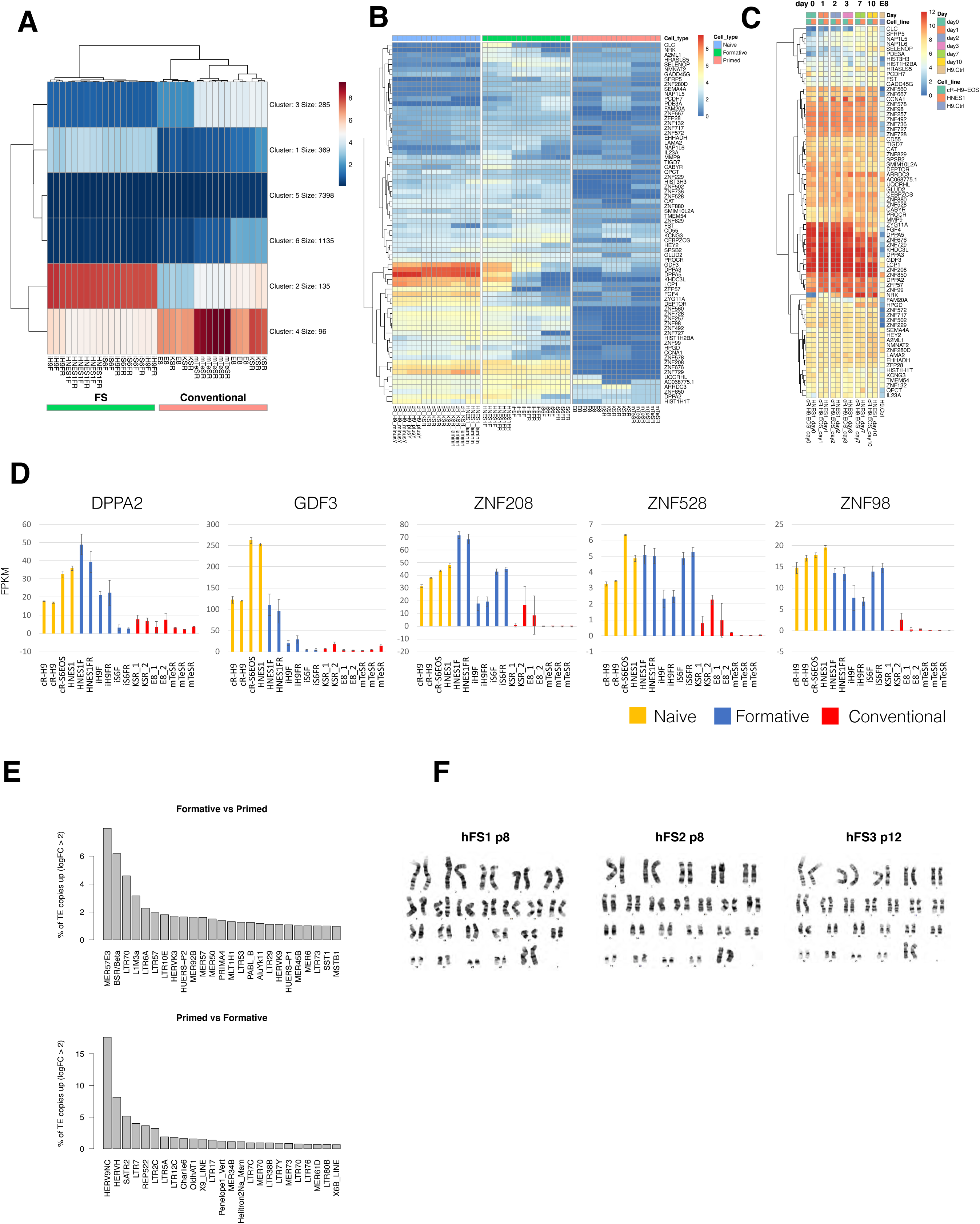
(A) K-means clustering of differentially expressed genes between human FS cells and conventional PSCs. (B) Gene expression heatmap of cluster 1 genes. Only protein coding genes (74 genes) are shown. (C) Expression heatmap of cluster 1 protein coding genes during naïve cell capacitation process (data from Rostovskaya et al 2019). (D) Related to Fig. 7I. FPKM values for additional selected naïve-formative specific genes. (E) Bar charts of differentially expressed TE families between FS cells and conventional hPSCs. (F) G-banding of chromosomes from three embryo derived hFS cell lines. 20 out of 20 metaphases were 46 (XX) for each line.

## Methods

### CONTACT FOR REAGENT AND RESOURCE SHARING

Further information and requests for resources and reagents should be directed to and will be fulfilled by the Lead Contact, Austin Smith.

### EXPERIMENTAL MODEL AND SUBJECT DETAILS

#### Ethics

Use of supernumerary human embryos in this research is approved by the Multi-Centre Research Ethics Committee, approval O4/MRE03/44, and licensed by the Human Embryology & Fertilisation Authority of the United Kingdom, research license R0178.

Experimental procedures using mice were carried out in facilities designated by the UK Home Office and are approved by the University of Cambridge Animal Welfare and Ethical Review Board and licensed by UK Home Office Project Licence 76777883.

## METHODS DETAILS

### Mouse FS cell, EpiSC and ES cell culture

FS cells were cultured in A_lo_XR medium, which consists of 3 ng/ml of activin A and 2 µM XAV939 and 1.0µM BMS439 in N2B27 medium (Nichols and Ying, 2006). EpiSCs were cultured in either AF (20ng/ml activin A and 12.5ng/ml Fgf2) or AFX (20ng/ml activin A, 12.5ng/ml Fgf2 and 2µM XAV939) in N2B27 medium. When passaging, cells were dissociated by Accutase into clumps and re-plated every 2-3 days at a ratio of 1:10-1:20. mES cells were maintained in 2i/LIF medium as described (Mulas et al., 2019). FS cells and EpiSCs were maintained under 7% CO_2_ and 5% O_2_ at 37°C on fibronectin (Fn) coated (16.7 µg/ml) plates and ES cells were maintained under 7% CO_2_ and 20% O_2_ at 37°C on 0.1 % gelatin coated plates. Experiments were generally performed between p10 and p30.

### Derivation of FS and EpiSCs from mouse embryo

E5.5 mouse embryos were dissected from decidua and further micro-dissected into embryonic and extraembryonic parts. Extra-embryonic endoderm layers were removed by mouth pipette and individual epiblasts were plated onto Fn coated (16.7 µg/ml) 4-well plates in either FS or EpiSC medium. After the epiblast outgrowth became large enough, the outgrowth was briefly incubated in Accutase and collected in wash buffer and re-plated onto a fresh 4-well plate.

### Derivation of FS and EpiSCs from mouse ES cells

ES cells were plated either directly in A_lo_XR, AF or AFX medium or N2B27 basal medium for two days and then re-plated in A_lo_XR, AF or AFX medium. Cultures were passaged at higher densities for the first 4-5 passages with Accutase.

### Derivation of human FS cells from Naïve PSCs

Naïve hES cells were differentiated in N2B27 medium for 7 days before changing to A_lo_XR. Cells were passaged every 3-5 days at a ratio of 1:10-1:20 and Rock inhibitor was added for the first 24 hours after dissociation. hFS cells were cultured on plates pre-coated with Laminin (10 µg/ml) and Fn (16.7 µg/ml).

### Derivation of human FS cell from embryos

Day 5 or day 6 human embryos, donated with informed consent from IVF programmes, were thawed using SAGE REF ART 8030 vitrification warming kit as per the manufacturer’s instructions and cultured for one or two days in N2B27 basal medium in 7% CO_2_ and 5% O_2_ at 37°C. ICMs were isolated on the following day by immunosurgery (Solter and Knowles, 1975) or mechanical dissociation and plated in A_lo_XR in the presence of Rock inhibitor on laminin/Fn coated 4-well plates. 2-4 weeks later, outgrowths were mechanically dissociated into clumps and replated into a fresh well. After this initial passage, Accutase was used for routine passaging.

### Embryoid body differentiation

2,000 cells were plated in low-binding 96-well plates in GMEM supplemented with 10% fetal calf serum, 2 mM L-glutamine, 0.1 mM Non-essential Amino Acid (NEAA) (GIBCO), 1 mM Sodium Pyruvate and 0.1 mM 2-ME. After 5 days, the EBs were transferred onto gelatin coated plates in fresh medium.

### PGCLC differentiation

3,000 mFS cells were plated in low-biding 96-well plates in GK15 medium (GMEM and 15 % Knockout Serum Replacement (GIBCO), 0.1 mM NEAA (GIBCO), 1 mM Sodium Pyruvate, 2 mM L-Glutamine, 0.1 mM 2-mercaptoethanol) supplemented with 500 ng/ml BMP2, 100ng/ml mSCF, 1 µg/ml hLIF, 50ng/ml Egf in the presence of 10 µM Rock inhibitor.

### Mesoderm differentiation

mFS cells were plated with 20 ng/ml activin a and 3 µM Chiron in N2B27 for 48 hours on Fn coated plate. hFS cells were plated with 3 µM CHIR99021 and 500 nM LDN193189 for the first 2 days followed by the addition of 20 ng/ml of Fgf2 from day3 to day6.

### Endoderm differentiation

mFS cells were plated with 20 ng/ml activin A and 3 µM Chiron in N2B27 for 24 hours and the medium was replaced thereafter with 20 ng/ml of activin A only for a further 2 days on Fn coated plate. hFS cells were differentiated in 100 ng/ml activin A, 100 nM PI-103, 3 µM CHIR99021, 10 ng/ml Fgf2, 3 ng/ml BMP4 and 10 µg/ml Heparin for the first 24hrs and then replaced with 100 ng/ml activin A, 100 nM PI-103, 20 ng/ml Fgf2, 250 nM LDN193189 and 10 µg/ml Heparin for a further 2 days.

### Neural differentiation

mFS cells were plated on laminin coated plate in N2B27 (Nichols and Ying, 2006). hFS cells were plated with 1 µM A83-01 and 500 nM LDN193189.

### Signal responsiveness experiment

Cells were plated in self-renewal medium and cultured overnight. On the following day, medium was changed to N2B27 medium with or without growth factors/inhibitors. The concentrations used were, activin A (20 ng/ml), Fgf2 (12.5 ng/ml), CHIR99021 (3 µM), Bmp2 (10 ng/ml), XAV939 (2 µM).

### Flow cytometry analysis

Mouse endoderm and mesoderm cells were dissociated with Cell Dissociation Buffer (GIBCO). mPGCLC were dissociated with TripLE Express (GIBCO). After the dissociation, cells were incubated with fluorophore conjugated antibodies in rat serum on ice for 20 min. Cells were washed once with wash buffer and analysed in HANK’s buffer supplemented with 1 % BSA. Antibodies are listed in key resource table.

### RT-qPCR

Total RNAs were purified by Reliaprep RNA miniprep kit (Promega). cDNAs were prepared by GoScript reverse transcription system (Promega). PCR was performed by Taqman Gene Expression Master Mix (Thermo Fisher Scientific) with Taqman (Thermo Fisher Scientific) or Universal Probe Library (Roche) probes. Probes and primer information are listed in Table S3.

### Immunofluorescence analysis

Cells were fixed on plates in 4% PFA for 15 minutes at RT. Cell were blocked with 5% skimmed milk or BSA/PBS 0.1 % TritonX. Primary and secondary antibodies were incubated for 1 hour at RT or overnight at 4°C. Antibodies used were listed in key resource table. Cells were imaged by LeicaDMI4000. PGCLCs and embryo sections were imaged by Leica SP5.

### FISH for *Xist*

FS cells were plated on Fn coated glass slide (Roboz Surgical instrument). The fluorescent conjugated RNA probe was purchased from Stellaris (Biosearch Technologies). *Xist* FSIH was performed as described previously (Sousa et al., 2018). Nuclear was stained with Dapi and imaged by Eclipse Ti Spinning Disk confocal microscope (Nikon).

### Metaphase chromosome analysis

FS Cells were treated with KaryoMAX colcemid (Gibco) and cultured further 2.5 hours. Cells were washed with PBS and harvested by Accutase and collected in wash buffer. After centrifuge, cells were resuspended in 5 ml of pre-warmed 0.075M KCl and incubated for 15 minutes at RT. Freshly prepared ice cold fixative solution (methanol: glacial acetic acid (3:1)) (100 µl) were added into the suspension and centrifuge. Cells were resuspended in 250-500 µl of fixative solution and up to 20 µl was spread onto a glass slide. DNA was counterstained with DAPI and spreads were imaged by Leica DMI4000 for counting. Karyotype analysis of embryo derived hFS cell lines were performed by Medical Genetics Service, Cytogenetics Laboratory, Cambridge University Hospitals.

### Immunoblotting

Culture plates were taken out from the incubator and placed on ice. Cells were washed with ice-cold PBS and lysed with RIPA buffer in the presence of Protease/Phosphatase inhibitor cocktail (Invitrogen). Lysed cells were rotated for 20 minutes and sonicated in Bioruptor (Diagnode). Cell lysates were cleared by centrifugation, and the supernatant was recovered. Protein concentrations were measured by the BCA method (Pierce). 25 µg of protein was loaded in each well. Blots were blocked with 5% BSA/TBS 0.1 % Triton-X for 1 hour at RT and incubated overnight with primary antibodies at 4°C. Secondary antibodies were incubated for 1 hour at RT and signals were detected with ECL Select (GE Healthcare) and Odyssey Fc (Li-Cor). 0.2 N NaOH was used when stripping.

### *Etv4/5* and *Otx2* knock out analysis

Etv4/5 dKO ES cell lines were established from Etv4 KO ES cells (Kalkan et al., 2019a) using a CRISPR/Cas9 based method. gRNAs were designed to excise Ets domain of Etv5 in Exon13 and Exon15. Otx2 KO ES cell lines were established from E14tg2a ES cells. gRNAs were designed to excise homeobox in Exon3. gRNAs were cloned into pCML32. Targeted ES cell clones were picked and genotyped by genomic PCR. Oct4 and Otx2 KO in FS cells were performed by co-transfected with one gRNA expression plasmid (pCML32, Oct4-1, Otx2-1 in Table S3, puromycin resistance, *piggybac* vector) with Cas9 expressing plasmid (G418 resistance, *piggybac* vector) and PBase expressing plasmid by TransIT LT1 (Mirus). Transfected cells were selected with 1 µg/ml of puromycin and 250 µg/ml of G418 from 24-48 hours post-transfection. Cells were counted and re-plated for another three days to form colonies. Rock inhibitor was added for the first 24 hours after replating. Alkaline phosphatase staining was performed following manufacture’s instruction (Sigma-Aldrich). gRNA sequences, genotyping primers and the amplicon sizes of each genotypes are listed in Table S3.

### RNA-sequencing

For the bulk RNA-sequencing experiment, cells were lysed in Trizol (Thermo Fisher Scientific) and total RNAs were prepared using the PureLink RNA Mini Kit (Thermo Fisher Scientific). Ribosomal RNAs were removed by Ribo-Zero rRNA Removal Kit (Illumina) and libraries were constructed using the NEXTflex Rapid Directional RNA-seq Kit (Bioo Scientific). For the low-input RNA-sequencing experiment, RNA was isolated from cells and epiblasts with the PicoPure RNA Isolation kit (Thermo Fisher Scientific) and libraries were constructed by SMARTerR Stranded Total RNA-Seq Kit v2 - Pico InputMammalian (Takara Clontech). 1,000 FS cells and entire isolated single epiblast from E5.0, E5.5, E6.0 were prepared per sample.

### ATAC-seq

50,000 cells were collected and washed with ice-cold PBS once then lysed in lysis buffer (10 mM Tris-HCl, pH 7.4, 10 mM NaCl, 3 mM MgCl_2_, 0.1% IGEPAL). The nuclear pellets were collected and Tn5 tagmentation and library construction performed using the Illumina Nextera kit (FC-121-1030). DNA was purified with AMPure XP beads (Beckman Coulter).

### ChIP-seq

ChIP was performed following the protocol reported previously (Kalkan et al., 2019a). Briefly, chromatin was cross-linked with 1% formaldehyde for 10 minutes at RT and quenched with 125 mM Glycine for 5 minutes at RT with rotation. After cell pellets were lysed, sonication was performed for 16 cycles on High setting, 30sec ON/30 sec OFF cycle by Bioruptor (Diagnode), 2×10^7^ cells per 300 µl in Bioruptor tube (Diagnode). 10% inputs were collected for the later library construction. Chromatin was immunoprecipitated with 2 µg of each antibodies and 20 µl of Protein G Dynabeads (Invitrogen) were used against 3×10^6^ cells. After the washes, DNA was eluted and each samples were treated with 2.5 µg/ml RNase A at 37°C for 30 minutes followed by 87.5 µg/ml Proteinase K at 55°C for 1 hour. DNA was purified with PCR clean-up kit (Qiagen). Libraries were prepared by NEXTflex Rapid DNA-Seq Kit 2.0 bundle with 96 HT barcodes (ParkinElmer).

### Single-cell RNA-seq

Cells were directly sorted into each well of 96-well plate filled with 2.3 µl of lysis buffer (1 unit/µl of SUPERaseIN RNase inhibitor (Invitrogen), 0.2 % Triton X) by BD FACSAria Fusion (BD Biosciences). Libraries were prepared using the Smart-seq2 protocol (Illumina) (Picelli et al., 2014).

### Chimaeras

#### FS cell chimaeras

FS cells were pre-treated with 10 µM Rock inhibitor for 1 hour before harvesting. Around 10 singly dissociated cells were injected into each blastocyst stage embryo. Embryos are either transferred into pseudo-pregnant mice or cultured *in vitro* for another 24 hours in N2B27. E9.5 mid-gestation stage embryos and juvenile mouse tissues were imaged by Leica stereo microscope. For sectioning, embryos and E12.5 gonads were replaced with 20% sucrose/PBS overnight at 4°C after the fixation then embedded in OCT compound and sectioned at 8 µm thickness. Sections were imaged by Zeiss apotome microscope or Leica SP5 confocal mocroscope.

#### Epiblast chimaeras

Homozygous mTmG mice were crossed with CD1 mice to obtain embryos. E5.5, 6.0-6.25 and E6.5 embryos were dissected from decidua and separated embryonic and extraembryonic halves. Extraembryonic endoderm layers were removed using a mouth-controlled pulled Pasteur pipette. Isolated epiblasts were treated with Accutase at room temperature and washed with M2 medium in the presence of 10 µM Rock inhibitor. Singly dissociated 10 cells were injected into blastocyst stage embryo of C57BL/6 mice. Micro-injection was performed in M2 medium with Rock inhibitor. For sectioning, embryo was embedded in OCT compound and sectioned in 10 µm thickness. Sections were stained with anti-RFP antibidies and imaged by Leica DMI4000.

### QUANTIFICATION AND STATISTICAL ANALYSIS

#### Bulk RNA-seq analysis

Low-quality RNA-seq reads and adaptor sequences were removed using *Trim Galore!*. Reads were aligned to the mouse (GRCm38/mm10) and human (GRCh38/hg38) reference genomes using *TopHat2* with parameters “--read-mismatch 2 --max-multihits 1 --b2-sensitive” considering uniquely mapping reads only. Gene counts were obtained using *featureCounts* using ENSEMBL (release 89) gene annotations. Normalization and differential expression analyses were performed using the R/Bioconductor *DESeq2* package. Normalized counts were transformed into log2 fragments per million (FPKM). Genes with log2 fold change>1.6 and adjusted p-value <0.05 were considered differentially expressed. Differentially expressed gene clusters for human cells were identified by k-means clustering of the first five principal components using the R ‘*kmeans*’ function. The distance plot was calculated using Euclidean distance between samples based on log2 normalized counts of expression values. Heatmaps were generated using the R ‘*pheatmap*” function.

For transposable elements (TEs), reads were aligned to the human (GRCh38/hg38) reference genome using *bowtie* with parameters “-a --best --strata -m 1 -v 2”, retaining uniquely mapping reads only in order to identify the genomic origin of TE transcription. Read counts on TEs were obtained using *featureCounts* on UCSC RepeatMasker-annotated regions. Normalization and differential expression analyses between cell types of identical genotype were performed with the R/Bioconductor *DESeq* package. TEs with an expression of at least log2-normalized counts > 3.5 in any cell type, a log2 fold change>2 and an adjusted p-value <0.05 were considered differentially expressed.

#### Published RNA-seq data comparison analysis

Mouse single cell RNA-seq data was downloaded from Nakamura et al., 2016 (GEO: GSE74767). Human naïve and conventional PSC transcriptome data were downloaded from SRA: SRP104789, ENA:E-MTAB-5114, ENA:E-MTAB-5674, GEO:GSE123005. The data was processed using the same methods as described above, except that genes with zero counts were removed from the single cell RNA-seq data matrix before further processing by DESeq2. The matrix of log2 fragment per millions for the *Macaca fascicularis* was obtained from GEO: GSE74767 (Nakamura et al., 2016). The Human single cell RNA-seq FPKM normalised counts matrix was downloaded from GEO: GSE136447 (Xiang et al., 2019).

#### PCA plots

Principal component analyses (PCA) were performed using the R ‘*prcomp*’ function based on log2-transformed Z-score expression values. To compare mouse and human bulk RNA-seq with mouse and macaque single cell RNA-seq, the principal components of the single cell RNA-seq data were calculated, with the bulk RNA-seq data projected onto this PCA space using the R ‘*predict’* function. These PCAs were computed using all expressed genes or with genes differentially expressed between the formative and primed lines in order to narrow down genes important for developmental progression. To compare human bulk RNA-seq with human single cell RNA-seq data, Log2 transformed counts were used. Using the most variable genes across the single cell stages, a PCA of the bulk samples was computed and the single cells were projected using the R ‘*predict*’ function.

#### scRNA-seq analysis

Raw files were quality controlled using FastQC v0.11.3 and results summarised with MultiQC, with checks including distributions of nucleotide content and sequencing depth. Reads were aligned to the *M*.*musculus* GRCm38.p6 reference genome with Ensembl v98 annotations using STAR v2.7.3a (--outSAMtype BAM SortedByCoordinate). Protein-coding gene quantification was done using Subread featureCounts v2.0.0 with Ensembl v98 annotations; only uniquely mapped reads were used. Cells with fewer than 3M reads were removed from further analysis, leaving 326 cells that passed the threshold. Raw expression levels were normalized using sctransform (Hafemeister and Satija, 2019), and the PCA created using the 2000 most abundant genes across the data. Jaccard similarity indices were calculated on the 2000 most abundant genes per cell, with similarities calculated between all cells of the same type.

#### GO-terms

Gene ontology (GO) term enrichment analyses were performed using the *David* tool.

#### ATAC-seq

Reads were quality-trimmed using *Trim Galore!*, and reads shorter than 15 nt were discarded. Reads were aligned to the mouse reference genome (GRCm38/mm10) using *bowtie* with parameters “-m1 -v1 --best --strata -X 2000 --trim3 1”. Duplicates were removed using *Picard tools*. Reads longer than one nucleosome length (146 nt) were discarded, and an offset of 4 nts was introduced. Peaks were called with *MACS2* and parameters “--nomodel --shift -55 --extsize 110 --broad -g mm --broad-cutoff 0.1”. Bigwig files for visualization on the UCSC Genome browser were generated using *deeptools bamcoverage* with parameters “–binSize 10 and --normalizeUsing RPKM”. ATAC peaks specific to each cell type were identified using *edgeR* within the R/Bioconductor *DiffBind* package using the option “bNot = T” to allow for contrasts between each cell type against all others. Significant peaks were determined using a log2 fold change of > 1 and FDR < 0.05. Heatmaps of ATAC-seq peaks were generated with *deeptools plotHeatmap*. DNA motif enrichment analyses for cell type-specific ATAC-seq peaks was performed using *HOMER*.

#### BS-seq

Whole genome BS-seq data was obtained from Zylicz et al., 2014 (GEO: GSE70355). BS-seq reads were aligned to the mouse reference genome (GRCm38/mm10) and deduplicated using *Bismark. MethPipe* was used calculate methylation levels at each CpG, and only CpGs with at least 5X read coverage were retained for further analyses. Methylation levels were averaged using a 250nt-sliding window to generate bigwig files.

#### ChIP-seq

Raw files were quality controlled using FastQC v0.11.3 and results summarised with MultiQC, with checks including distributions of nucleotide content, sequencing depth and adapter contamination. Reads were aligned to the *M*.*musculus* GRCm38.p6 reference genome using bwa mem v0.7.10-r789 (default parameters); the MT, X, Y chromosomes and scaffolds were excluded from the resulting BAM files. Genome browser tracks for the UCSC genome browser were created with deepTools bamCoverage v3.3.1 (—binSize 30). Averaged genome browser tracks for ChIP profile visualization were created as follows: first the tracks were generated with bamCoverage (—binSize 5 –normalizeUsing RPKM), then the output was averaged using wiggletools v1.2.1 (Zerbino et al., 2014). Profiles of the ChIP tracks on the ATAC peaks were created using deepTools computeMatrix (reference-point --binSize 5 -b 4000 -a 4000 --referencePoint center) and plotProfile (default parameters). To identify bivalent promoters, peak regions were called with macs2 v2.2.6 (-f BAMPE -q 0.05), only peaks with signalValue>5 were considered for downstream analysis. Peak regions were intersected per condition and across histone marks using bedops v2.4.38. HOMER v4.10 was used to calculate distance between peaks and transcription start sites (mm10 -size 3000); peaks within 3kb of a TSS were considered as promoter peaks.

## DATA AND SOFTWARE AVAILABILITY

Sequencing data were deposited to the GEO.

