## Supplemental figures and legends for "Capture of mouse and human stem cells with features of formative pluripotency"

**Figure S7, related to Figure 7**

(A) K-means clustering of differentially expressed genes between human FS cells and conventional PSCs. (B) Gene expression heatmap of cluster 1 genes. Only protein coding genes (74 genes) are shown. (C) Expression heatmap of cluster 1 protein coding genes during naïve cell capacitation process (data from Rostovskaya et al 2019). (D) Related to Fig. 7I. FPKM values for additional selected naïve-formative specific genes. (E) Bar charts of differentially expressed TE families between FS cells and conventional hPSCs. (F) G-banding of chromosomes from three embryo derived hFS cell lines. 20 out of 20 metaphases were 46 (XX) for each line.

#### FigS.1

A

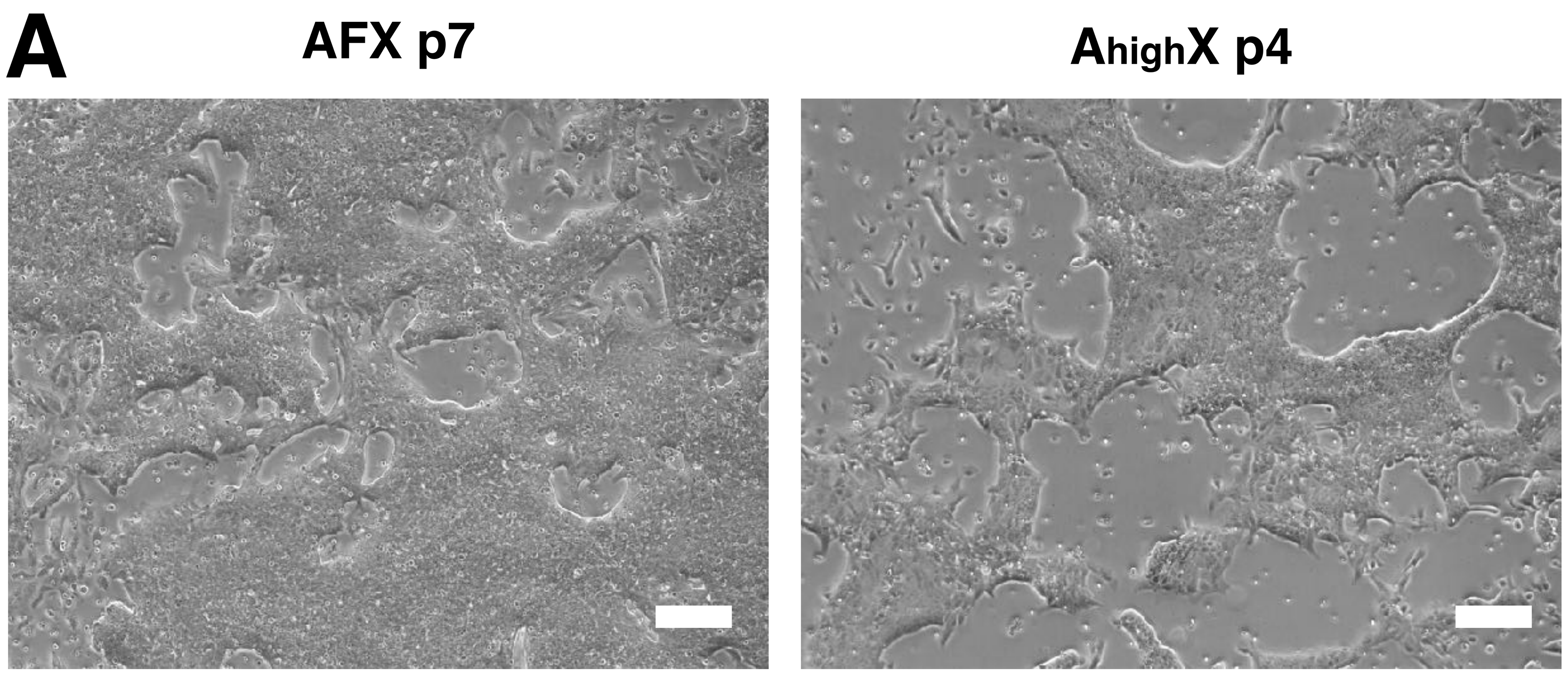


B

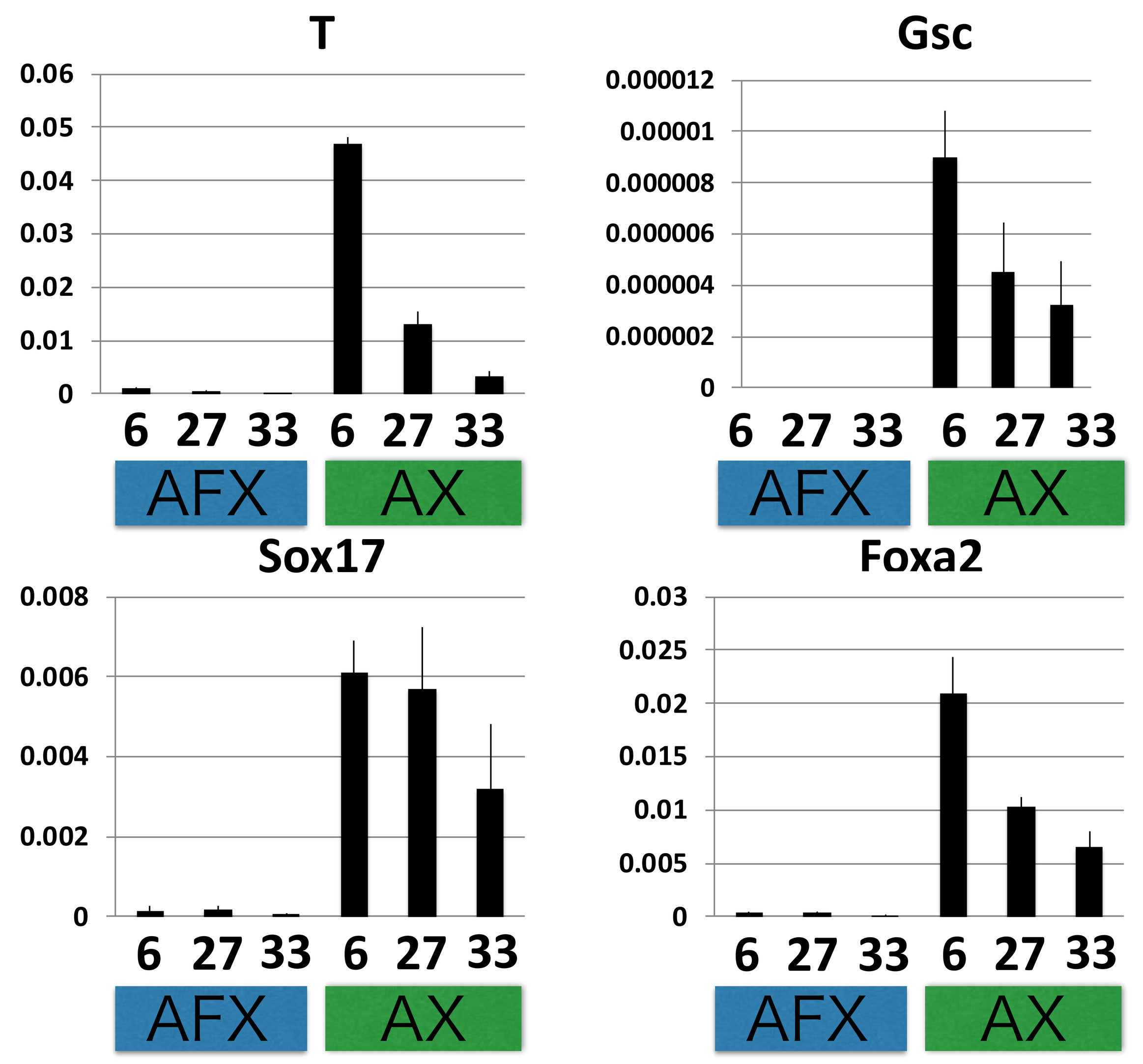


C

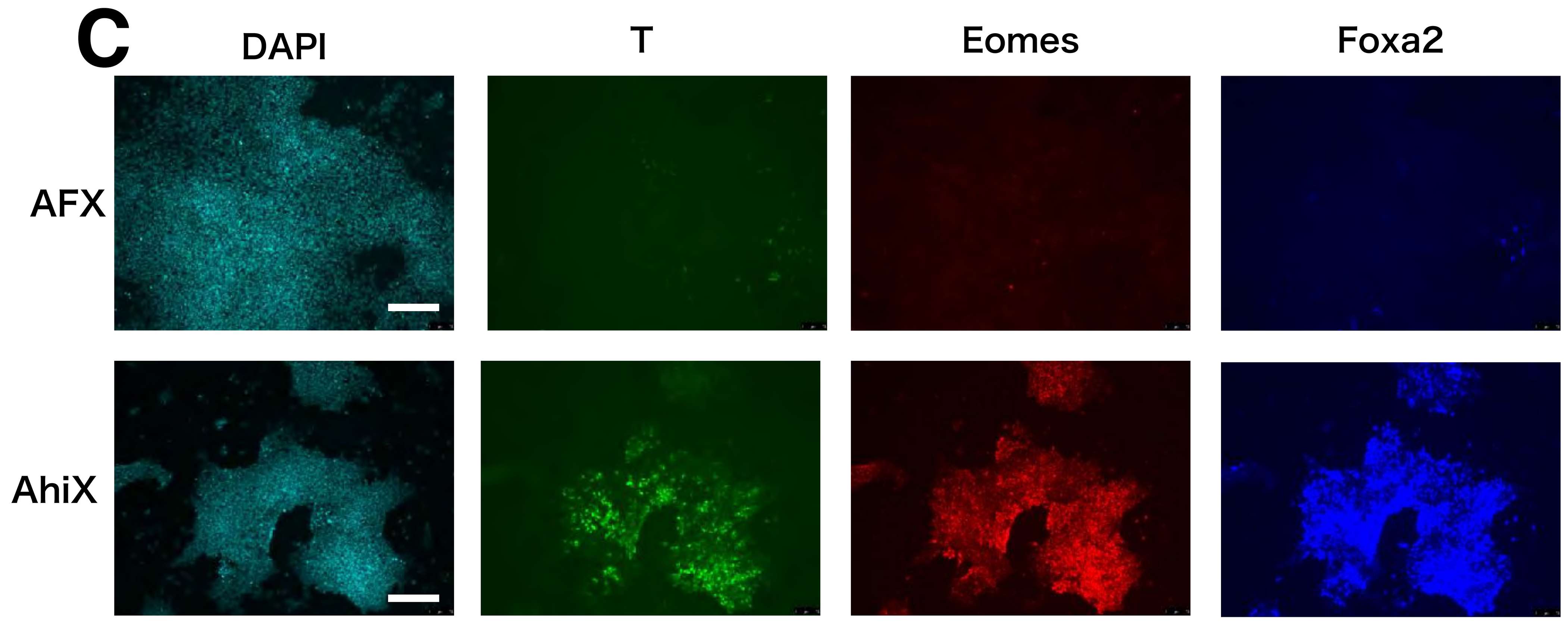


D

| Concentration of activin | No of E5.5 embryos | No of established lines (> p10) | Success rate |
|---|---|---|---|
| 1ng | 8 | 0 | 0.0% |
| 1.5ng | 8 | 1 | 12.5% |
| 2ng | 11 | 3 | 27.3% |
| 2.5ng | 4 | 0 | 0.0% |
| 3ng | 11 | 4 | 36.4% |

E

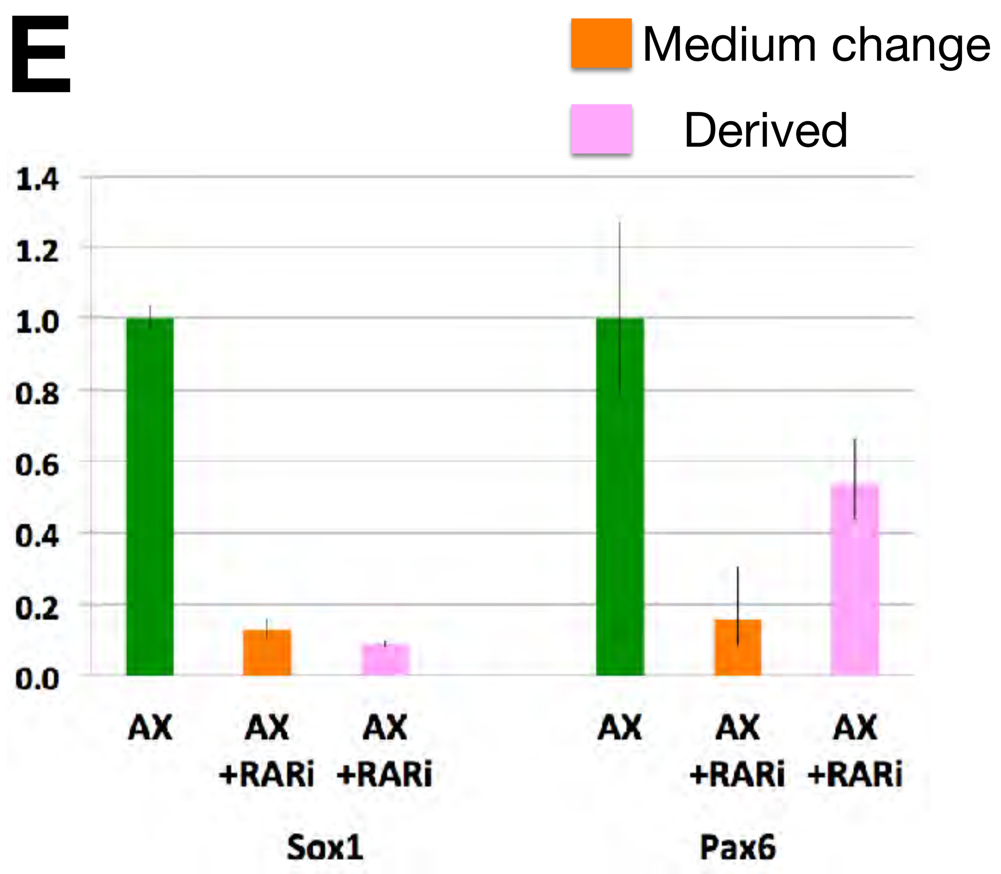


F

| | Number of E5.5 embryos | Lines established | Efficiency |
|---|---|---|---|
| AX | 8 | 4 | 50% |
| AX +RARi | 4 | 4 | 100% |

G

| Line | Percentage of diploid (=40) cells | Passage number |
|---|---|---|
| 5a6 | 75.0 % (24/32) | 19 |
| NBRA3.2 | 87.1 % (27/31) | 22 |
| 5ar1 | 64.0 % (16/25) | 21 |
| 5ar2 | 83.3% (20/24) | 26 |

H

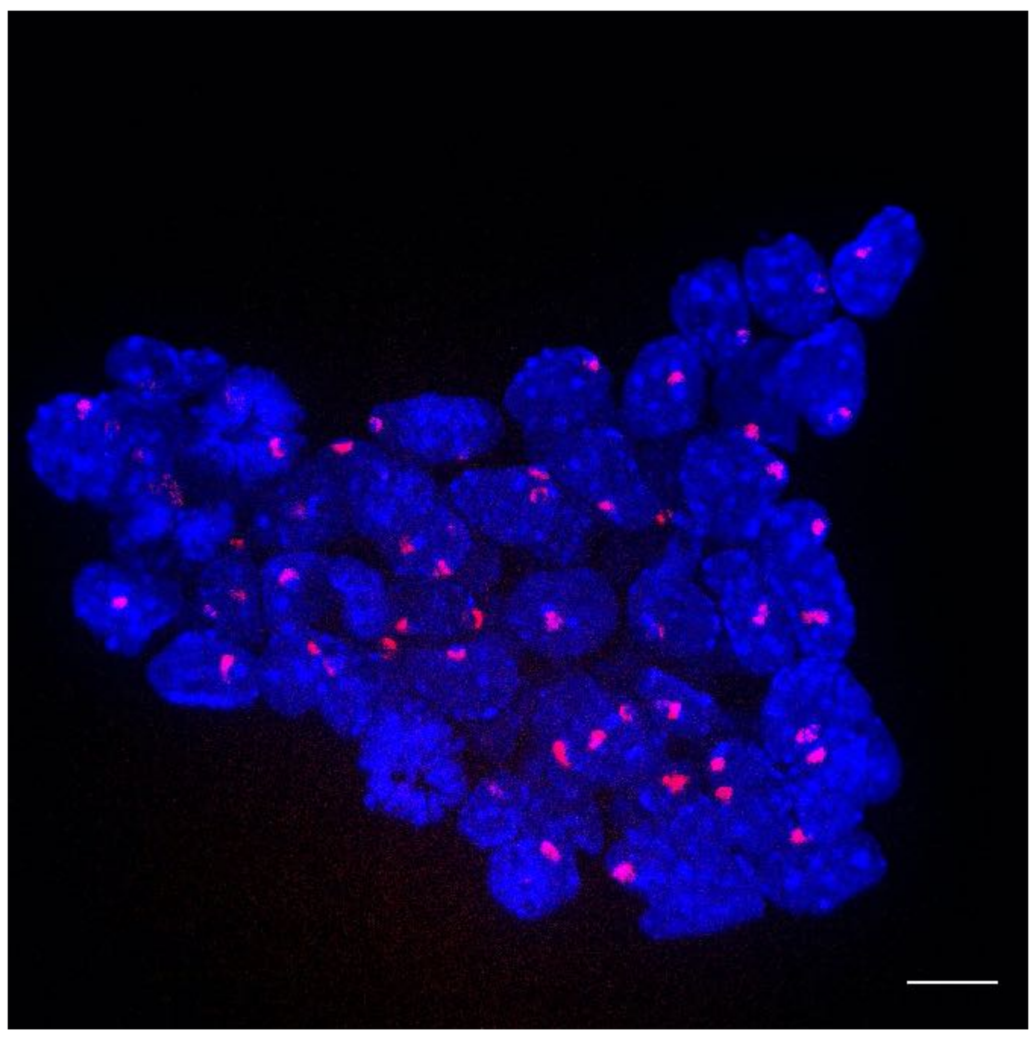

I

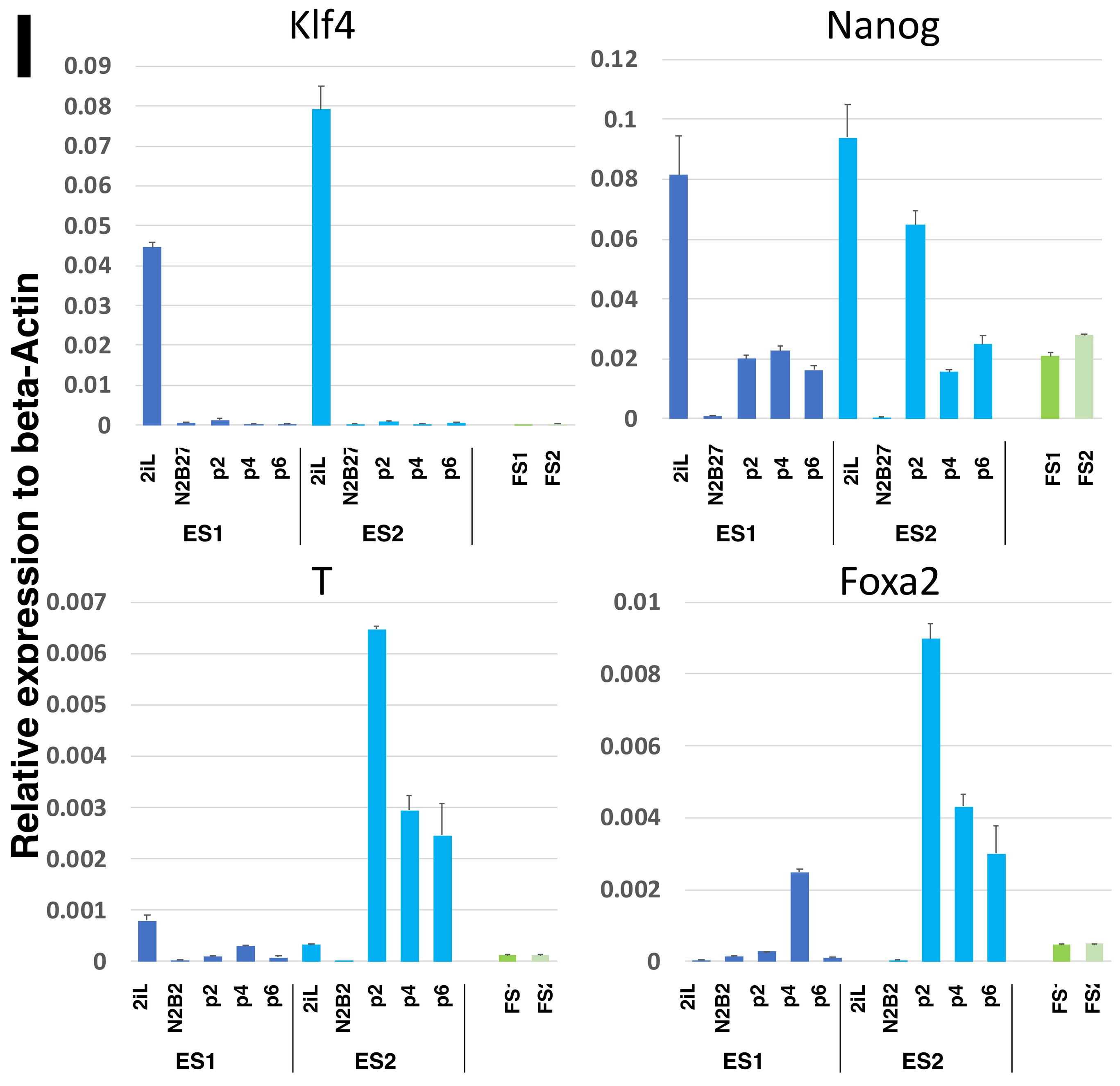


J

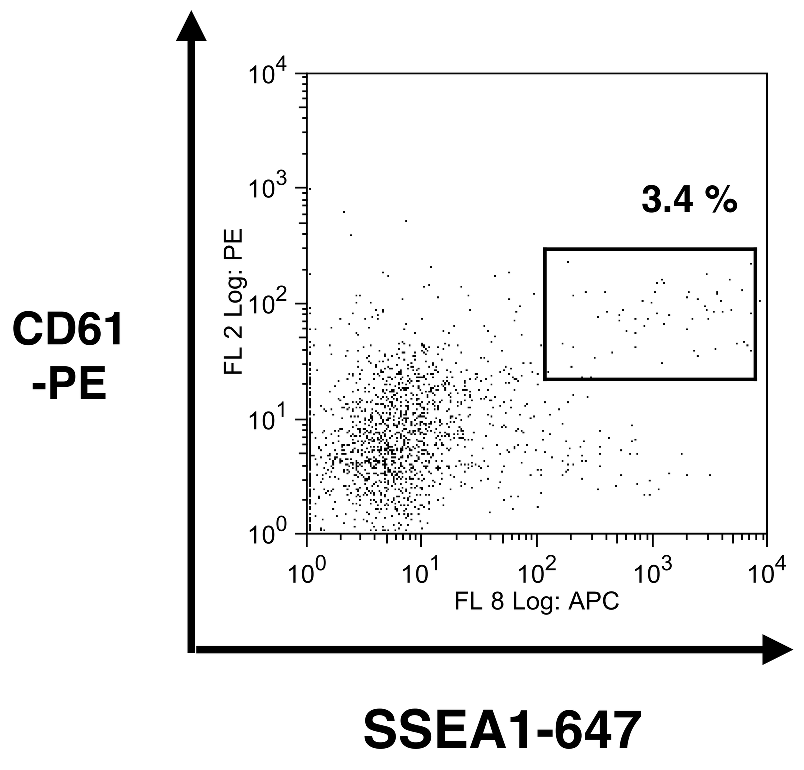


K

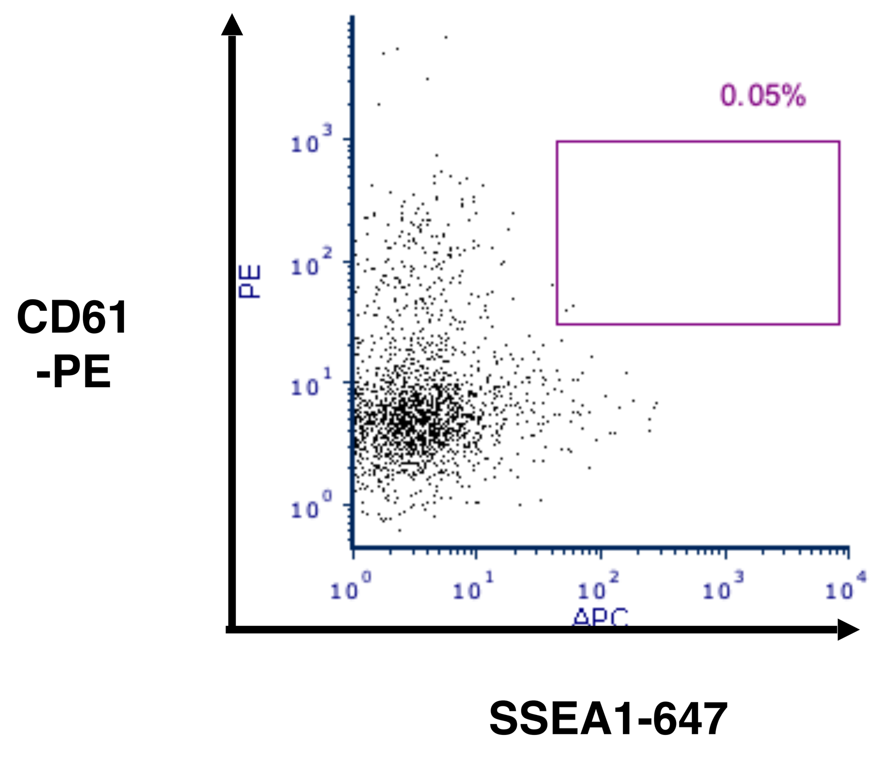


L

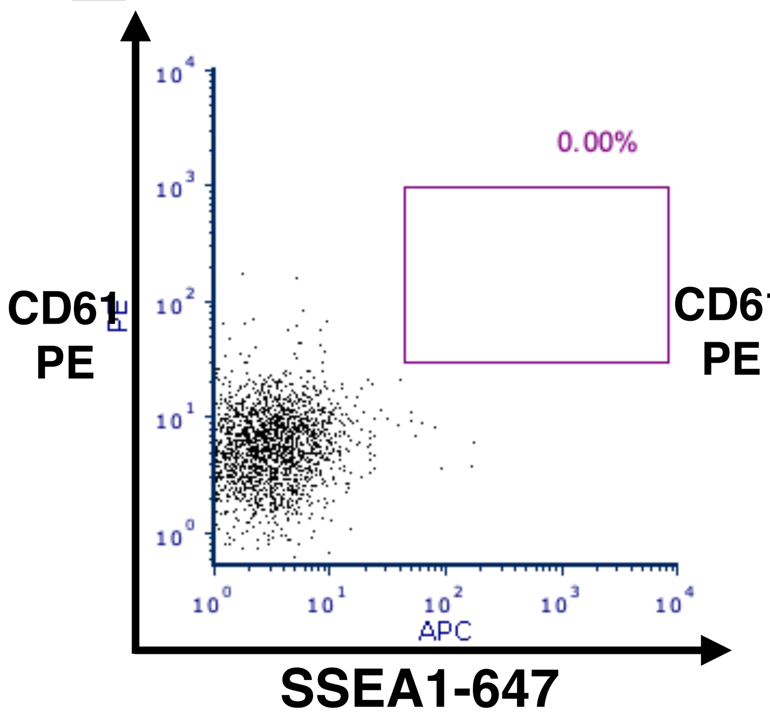


M

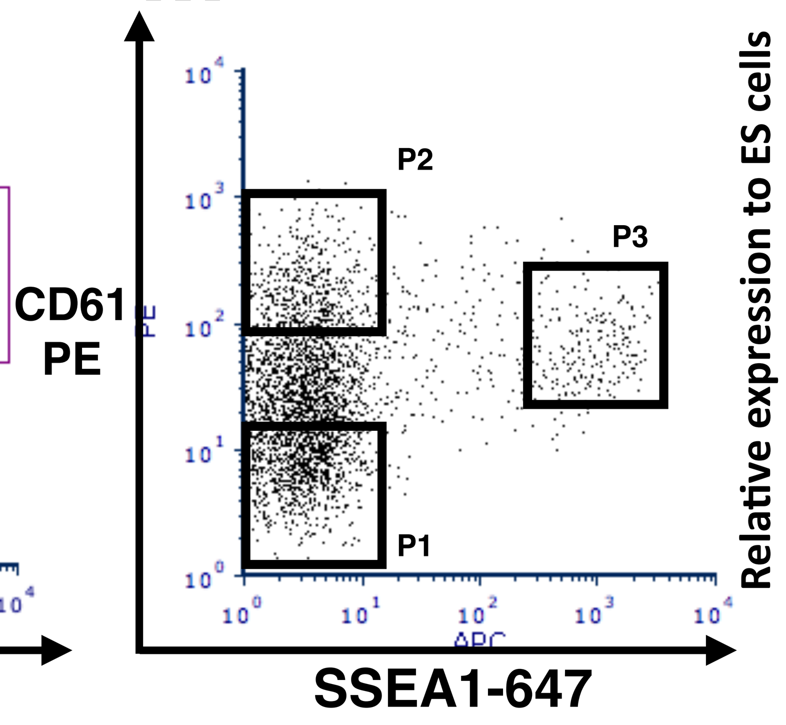


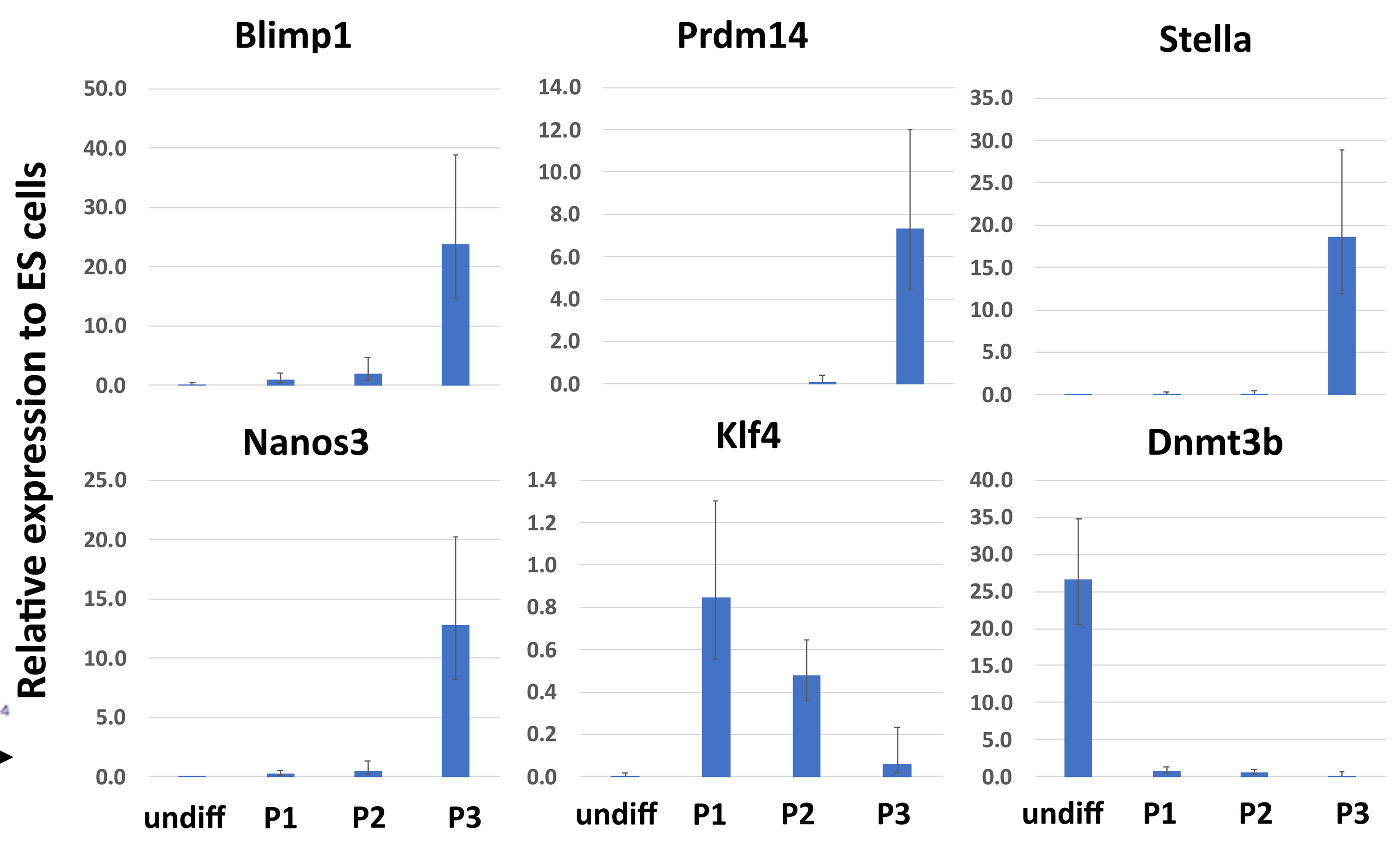


N

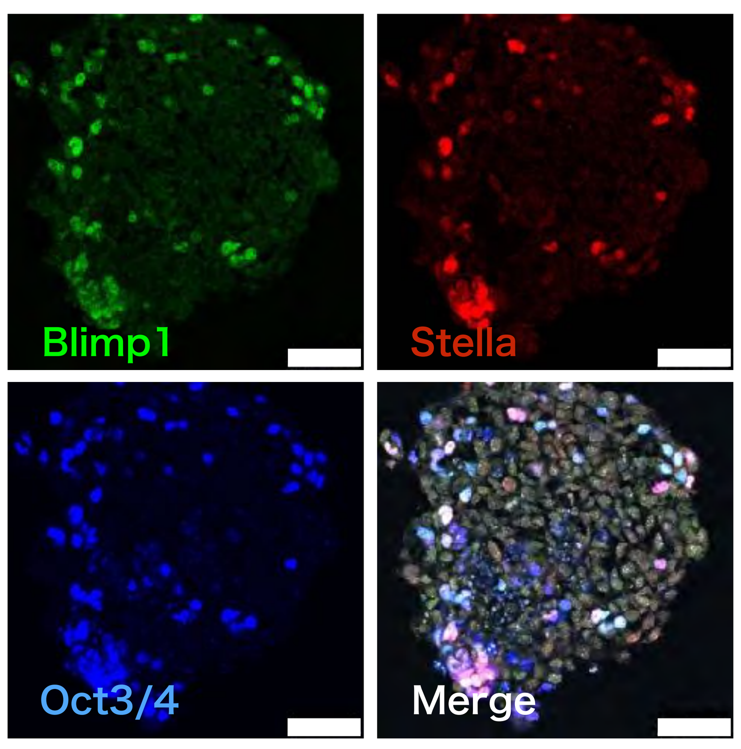

### FigS2

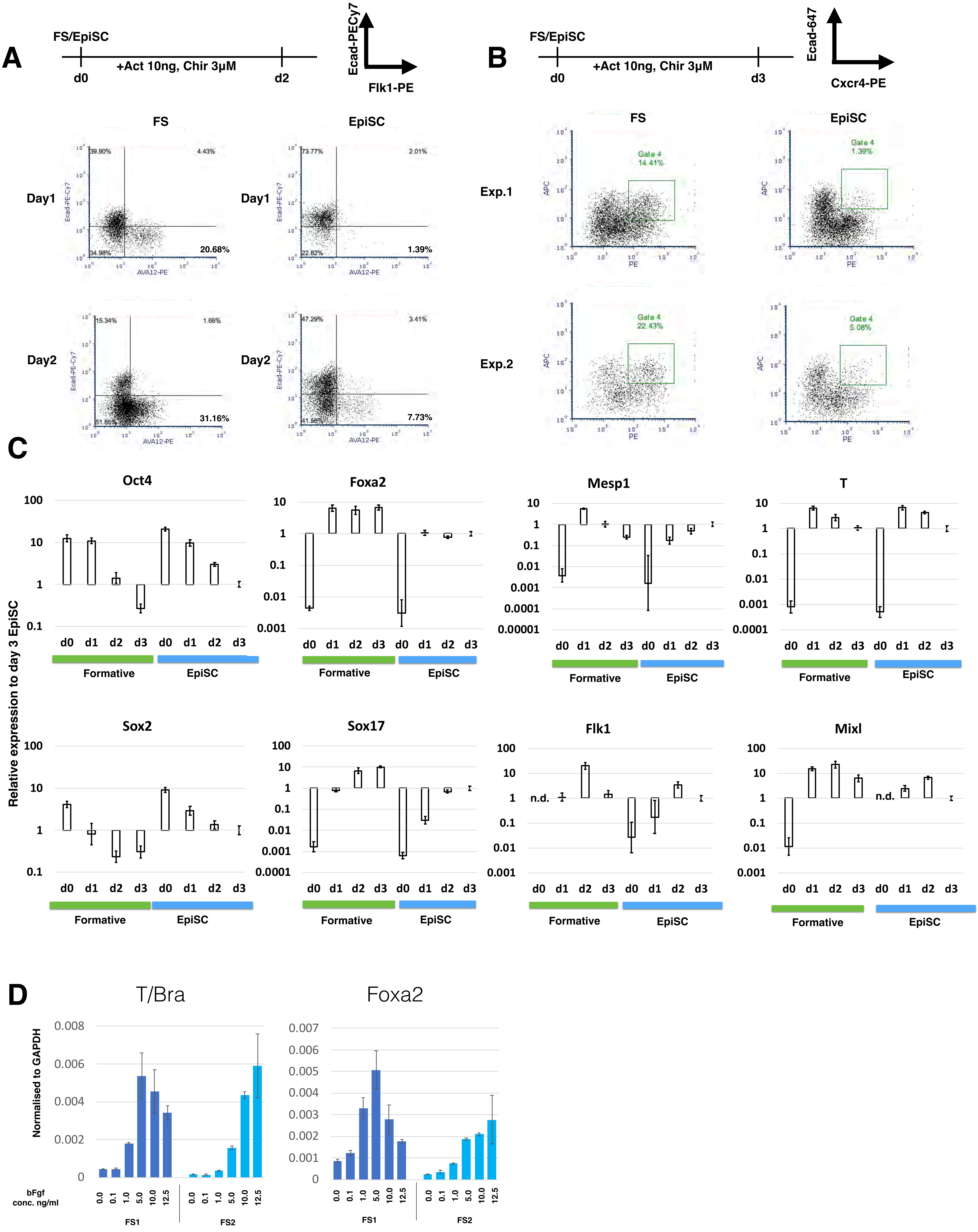

A
FS/EpiSC
+Act 10ng, Chir 3μM
d0
d2
Ecad-PECy7
Flk1-PE
FS
EpiSC
Day1
Day2
39.90%
4.43%
34.98%
20.68%
73.77%
2.01%
22.82%
1.39%
15.34%
1.66%
51.85%
31.16%
47.29%
3.41%
41.56%
7.73%
Ecad-PE-Cy7
AVA12-PE
B
FS/EpiSC
+Act 10ng, Chir 3μM
d0
d3
Ecad-647
Cxcr4-PE
FS
EpiSC
Exp.1
Exp.2
Gate 4
14.41%
1.39%
22.43%
5.08%
APC
PE
C
Relative expression to day 3 EpiSC
Oct4
Foxa2
Mesp1
T
Sox2
Sox17
Flk1
Mixl
n.d.
d0 d1 d2 d3 d0 d1 d2 d3
Formative
EpiSC
D
T/Bra
Foxa2
Normalised to GAPDH
bFgf conc. ng/ml
0.0 0.1 1.0 5.0 10.0 12.5
FS1
FS2

Fig S3

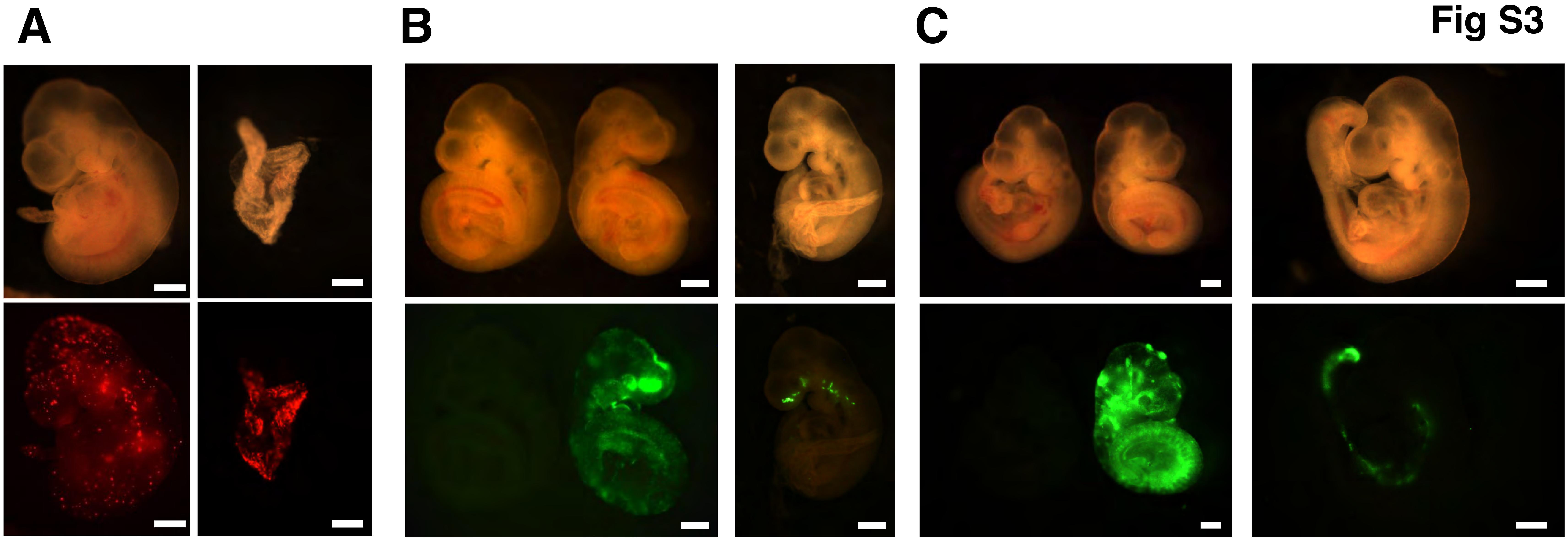


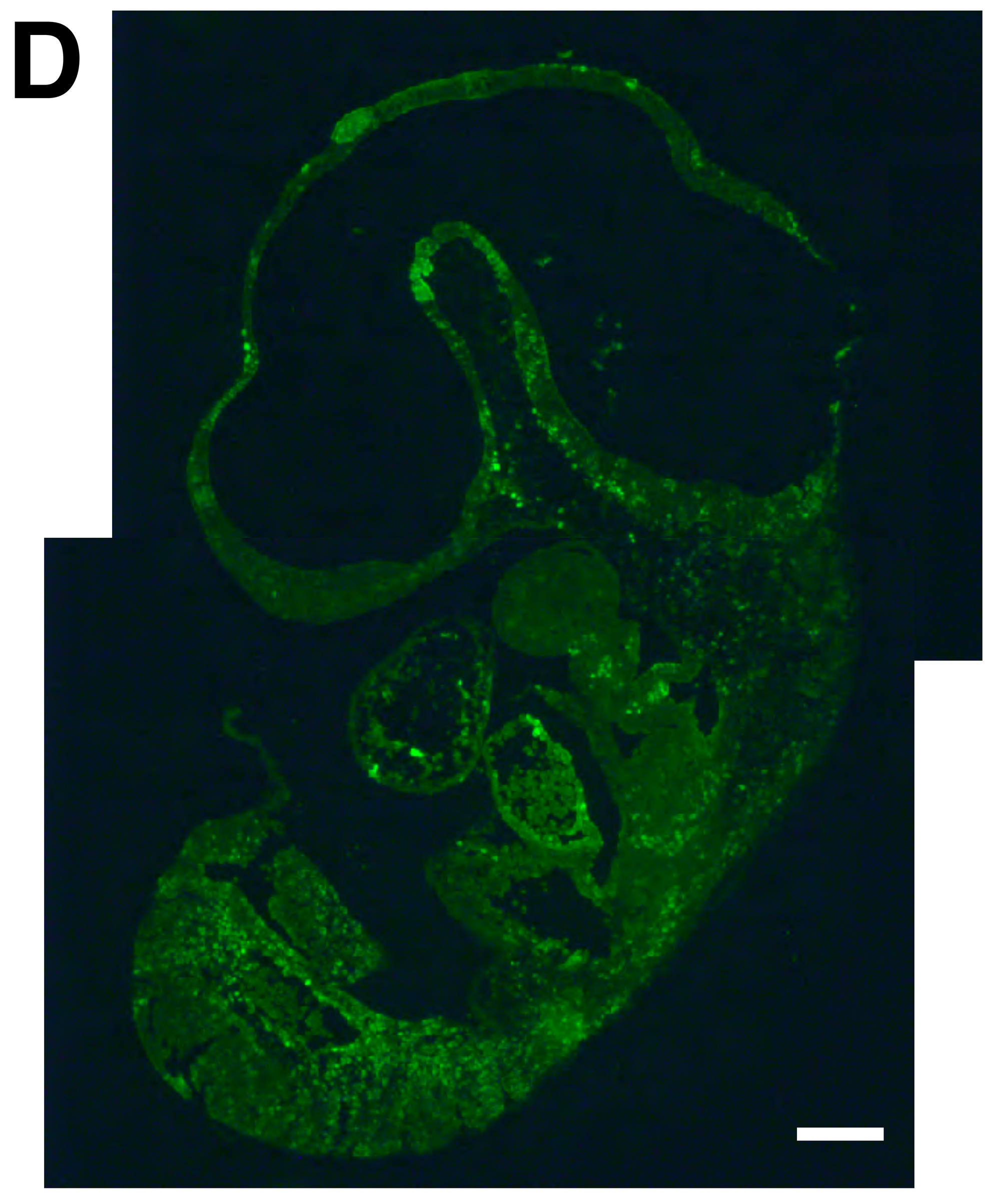


E

| Line | Embryos recovered | Widespread Contribution | Localised contribution | Yolk sac contribution | Chimaera rate |
|---|---|---|---|---|---|
| NBRA3.2 | 48 | 8 | 11 | 1* | 39.6% |
| 5a6 | 20 | 1 | 2 | 1 | 15.0% |
| 5ar1 | 17 | 1 | 3 | Not Investigated | 23.5% |

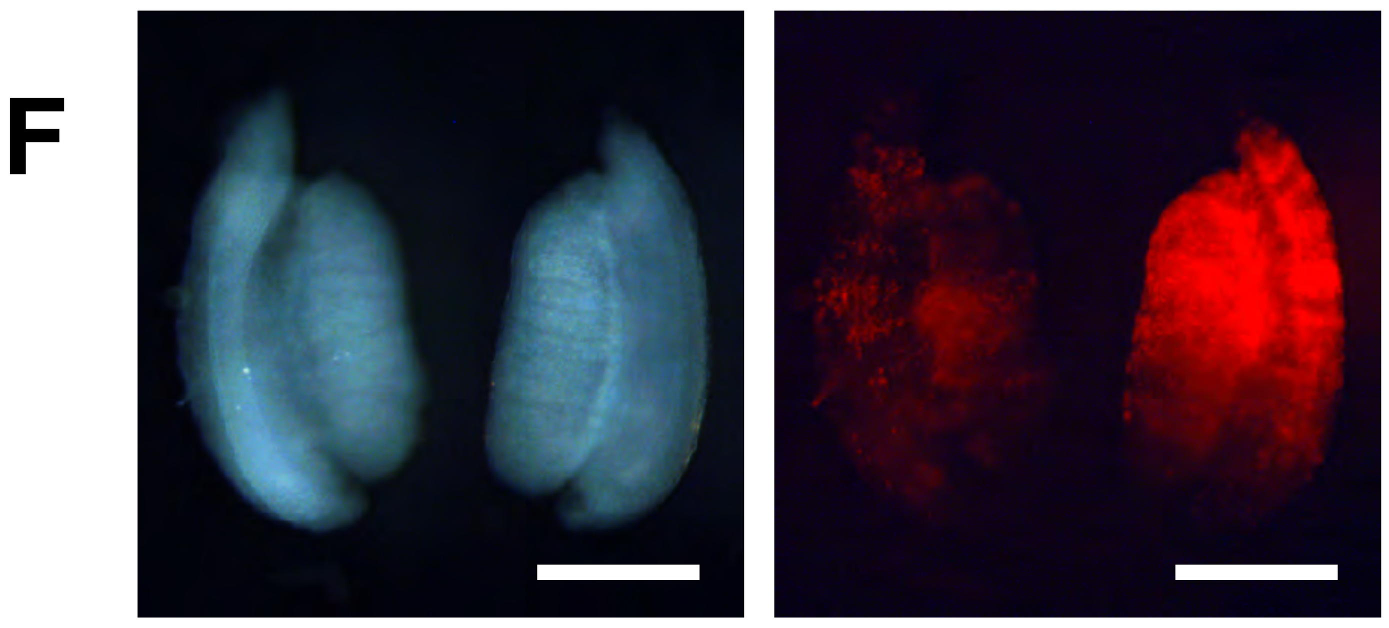


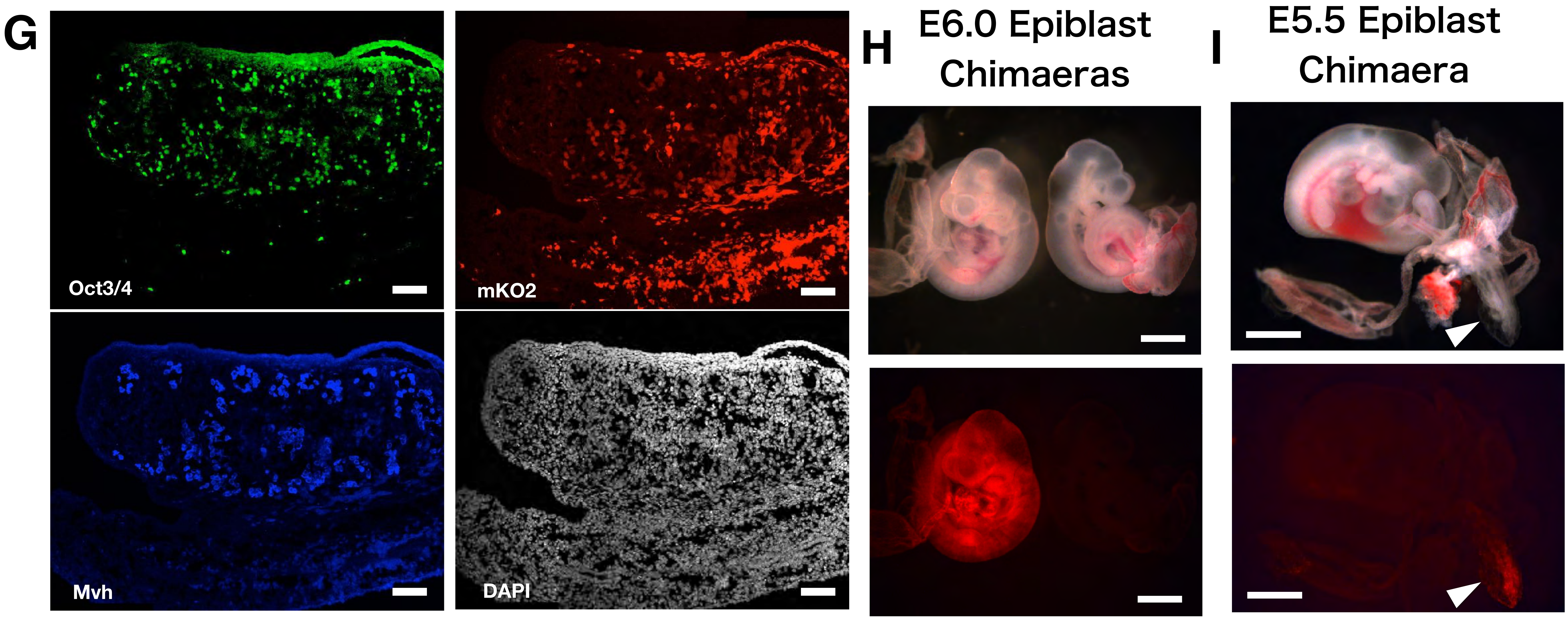


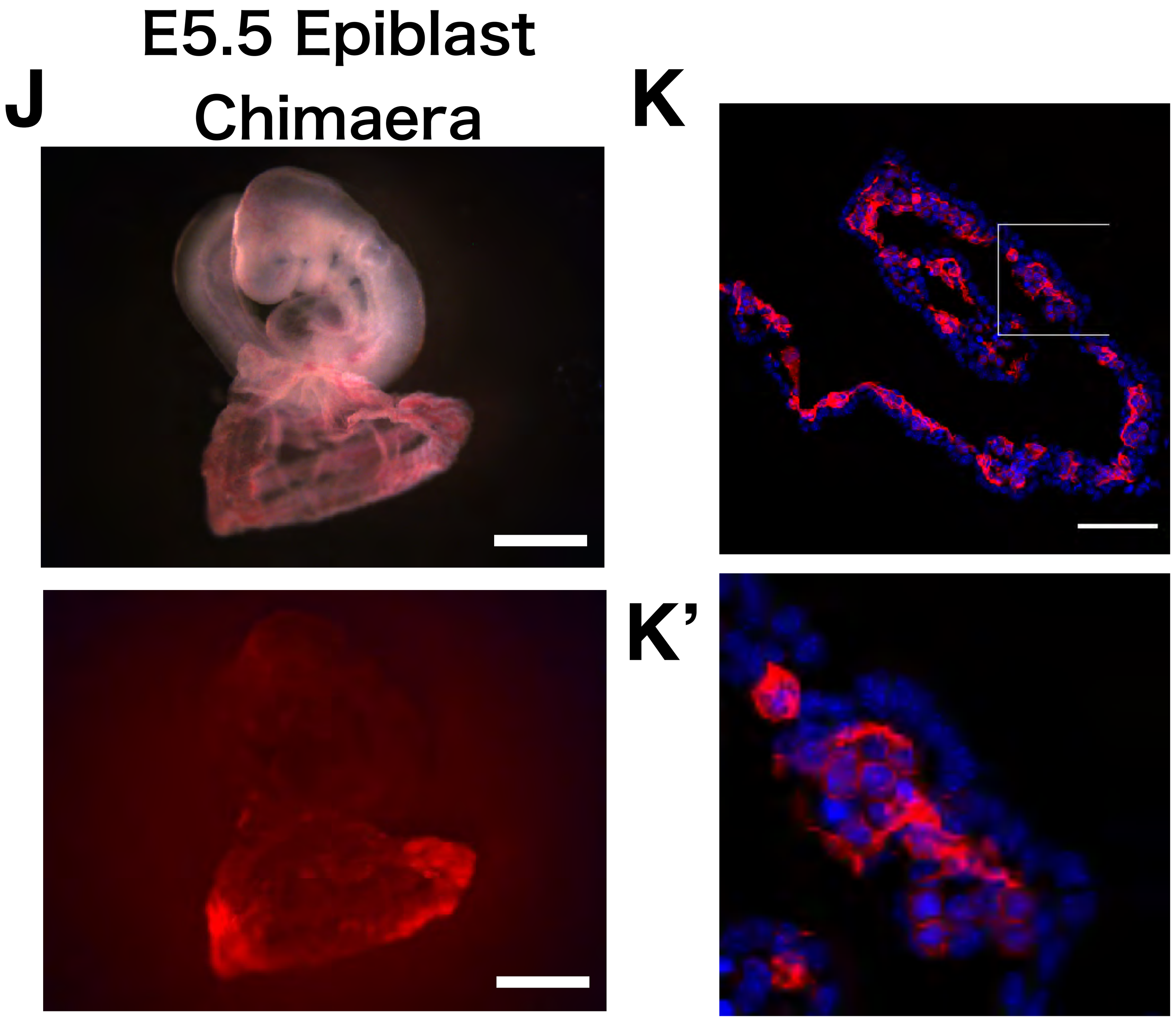


L

| Donor stage | Embryos recovered | Widespread Contribution | Localised contribution | contribution yolk sac and/or amnion | Chimaera rate |
|---|---|---|---|---|---|
| E5.5 | 44 | 2 | 5 | 7 | 16.0% |
| E6.0 | 47 | 1 | 3 | 4 | 8.5% |
| E6.5 | 29 | 0 | 0 | 0 | 0.0% |

FigS4

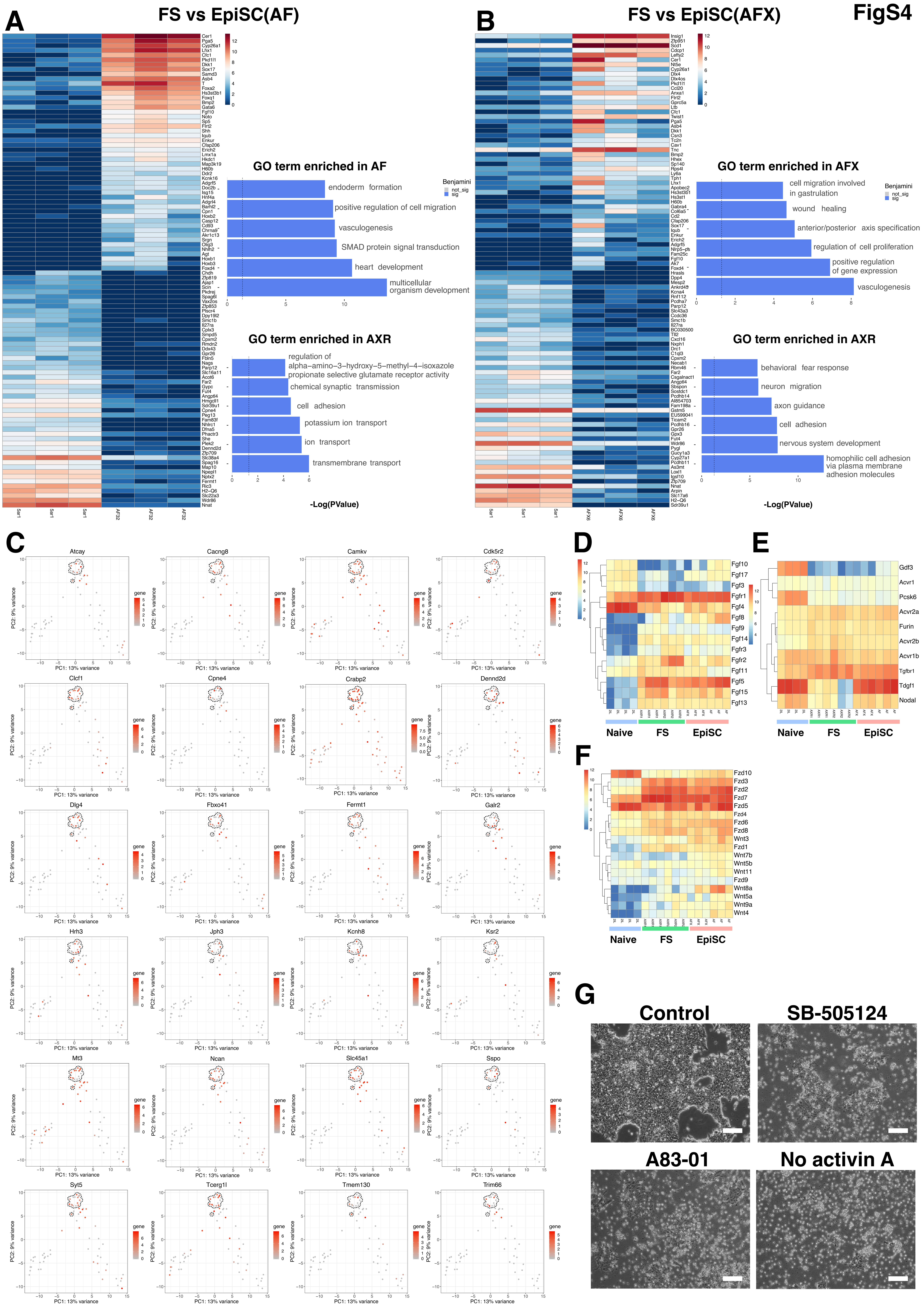
A
FS vs EpiSC(AF)
GO term enriched in AF
endoderm formation
positive regulation of cell migration
vasculogenesis
SMAD protein signal transduction
heart development
multicellular organism development
Benjamini
not_sig
sig
GO term enriched in AXR
regulation of alpha−amino−3−hydroxy−5−methyl−4−isoxazole propionate selective glutamate receptor activity
chemical synaptic transmission
cell adhesion
potassium ion transport
ion transport
transmembrane transport
-Log(PValue)
B
FS vs EpiSC(AFX)
GO term enriched in AFX
cell migration involved in gastrulation
wound healing
anterior/posterior axis specification
regulation of cell proliferation
positive regulation of gene expression
vasculogenesis
GO term enriched in AXR
behavioral fear response
neuron migration
axon guidance
cell adhesion
nervous system development
homophilic cell adhesion via plasma membrane adhesion molecules
-Log(PValue)
C
Atcay
Cacng8
Camkv
Cdk5r2
Clcf1
Cpne4
Crabp2
Dennd2d
Dlg4
Fbxo41
Fermt1
Galr2
Hrh3
Jph3
Kcnh8
Ksr2
Mt3
Ncan
Slc45a1
Sspo
Syt5
Tcerg1l
Tmem130
Trim66
PC1: 13% variance
PC2: 9% variance
gene
D
Fgf10
Fgf17
Fgf3
Fgfr1
Fgf4
Fgf8
Fgf9
Fgf14
Fgfr3
Fgfr2
Fgf11
Fgf5
Fgf15
Fgf13
Naive
FS
EpiSC
E
Gdf3
Acvr1
Pcsk6
Acvr2a
Furin
Acvr2b
Acvr1b
Tgfbr1
Tdgf1
Nodal
Naive
FS
EpiSC
F
Fzd10
Fzd3
Fzd2
Fzd7
Fzd5
Fzd4
Fzd6
Fzd8
Wnt3
Fzd1
Wnt7b
Wnt5b
Wnt11
Fzd9
Wnt8a
Wnt5a
Wnt9a
Wnt4
Naive
FS
EpiSC
G
Control
SB-505124
A83-01
No activin A

Fig.S5

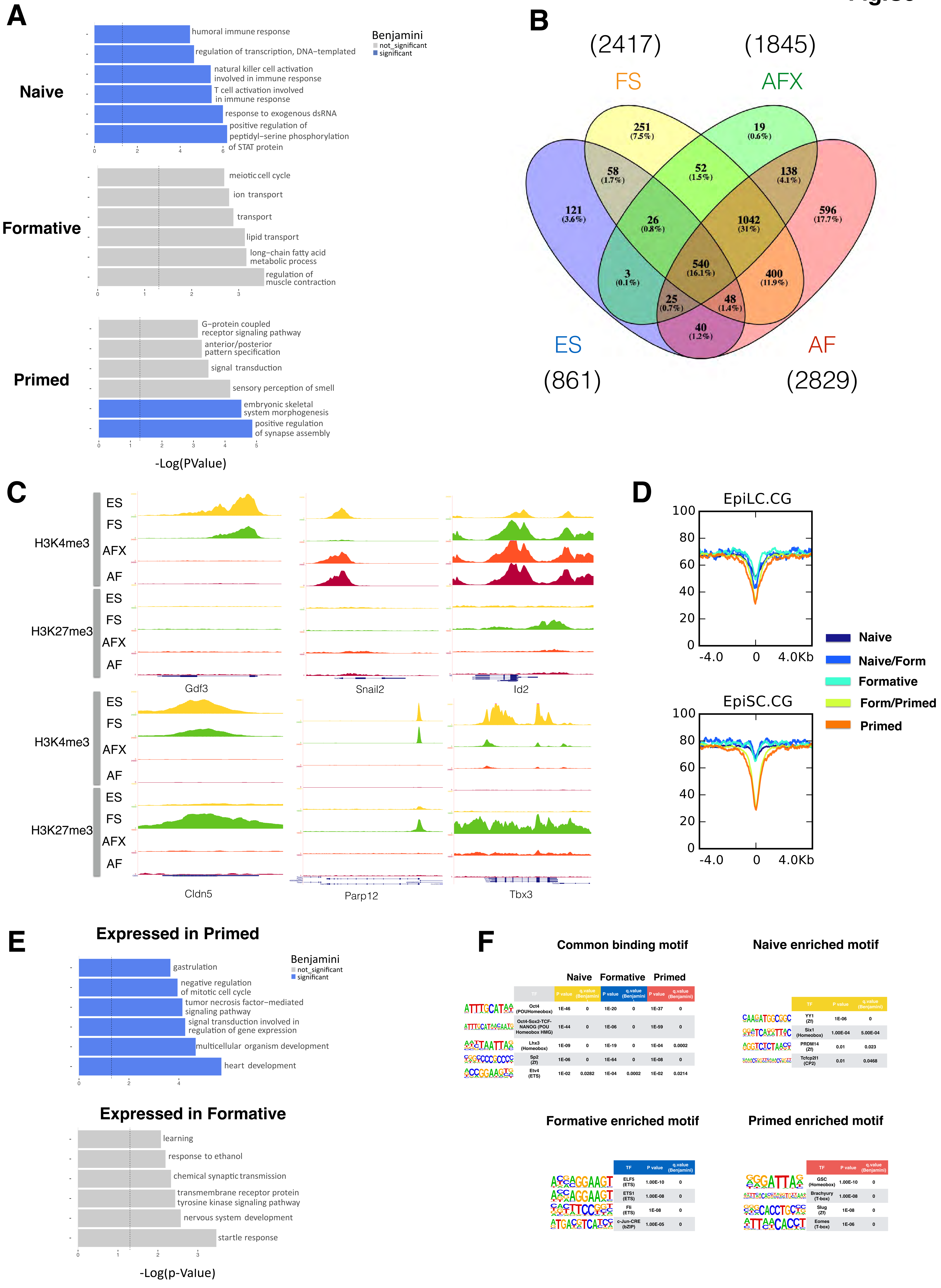
A
Naive
Formative
Primed
Benjamini
not_significant
significant
humoral immune response
regulation of transcription, DNA−templated
natural killer cell activation involved in immune response
T cell activation involved in immune response
response to exogenous dsRNA
positive regulation of peptidyl−serine phosphorylation of STAT protein
meiotic cell cycle
ion transport
transport
lipid transport
long−chain fatty acid metabolic process
regulation of muscle contraction
G−protein coupled receptor signaling pathway
anterior/posterior pattern specification
signal transduction
sensory perception of smell
embryonic skeletal system morphogenesis
positive regulation of synapse assembly
-Log(PValue)
B
(2417)
(1845)
FS
AFX
ES
AF
(861)
(2829)
C
H3K4me3
H3K27me3
ES
FS
AFX
AF
Gdf3
Snail2
Id2
Cldn5
Parp12
Tbx3
D
EpiLC.CG
EpiSC.CG
-4.0
0
4.0Kb
Naive
Naive/Form
Formative
Form/Primed
Primed
E
Expressed in Primed
gastrulation
negative regulation of mitotic cell cycle
tumor necrosis factor−mediated signaling pathway
signal transduction involved in regulation of gene expression
multicellular organism development
heart development
Expressed in Formative
learning
response to ethanol
chemical synaptic transmission
transmembrane receptor protein tyrosine kinase signaling pathway
nervous system development
startle response
-Log(p-Value)
F
Common binding motif
Naive enriched motif
Naive
Formative
Primed
Formative enriched motif
Primed enriched motif

Fig. S6

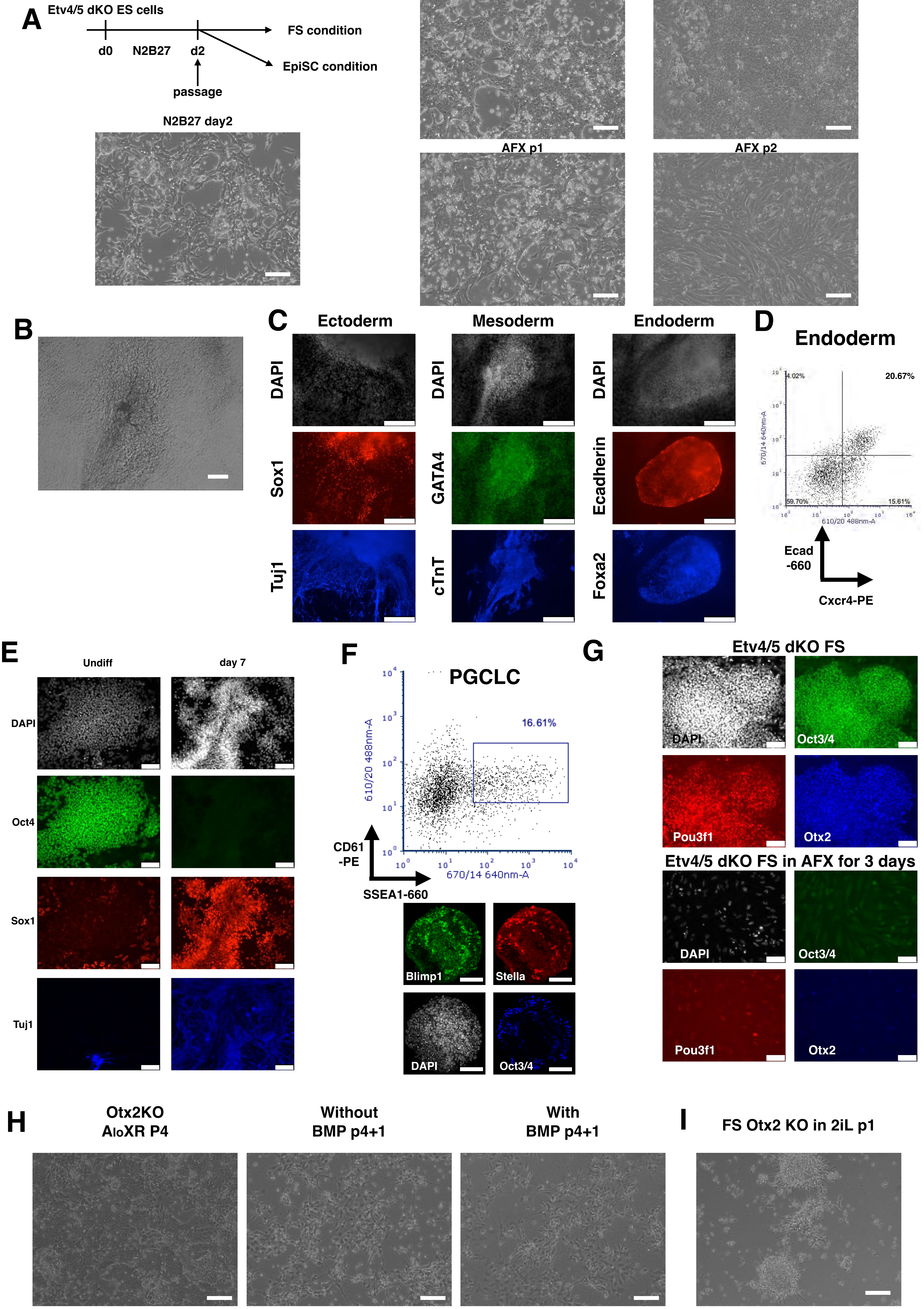
A
Etv4/5 dKO ES cells
d0
N2B27
d2
passage
FS condition
EpiSC condition
N2B27 day2
FS p1
FS p2
AFX p1
AFX p2
B
C
Ectoderm
Mesoderm
Endoderm
DAPI
Sox1
Tuj1
GATA4
cTnT
Ecadherin
Foxa2
D
Endoderm
4.02%
20.67%
59.70%
15.61%
670/14 640nm-A
610/20 488nm-A
Ecad
-660
Cxcr4-PE
E
Undiff
day 7
DAPI
Oct4
Sox1
Tuj1
F
PGCLC
16.61%
610/20 488nm-A
670/14 640nm-A
CD61
-PE
SSEA1-660
Blimp1
Stella
DAPI
Oct3/4
G
Etv4/5 dKO FS
DAPI
Oct3/4
Pou3f1
Otx2
Etv4/5 dKO FS in AFX for 3 days
DAPI
Oct3/4
Pou3f1
Otx2
H
Otx2KO
AloXR P4
Without
BMP p4+1
With
BMP p4+1
I
FS Otx2 KO in 2iL p1

**Fig. S7**

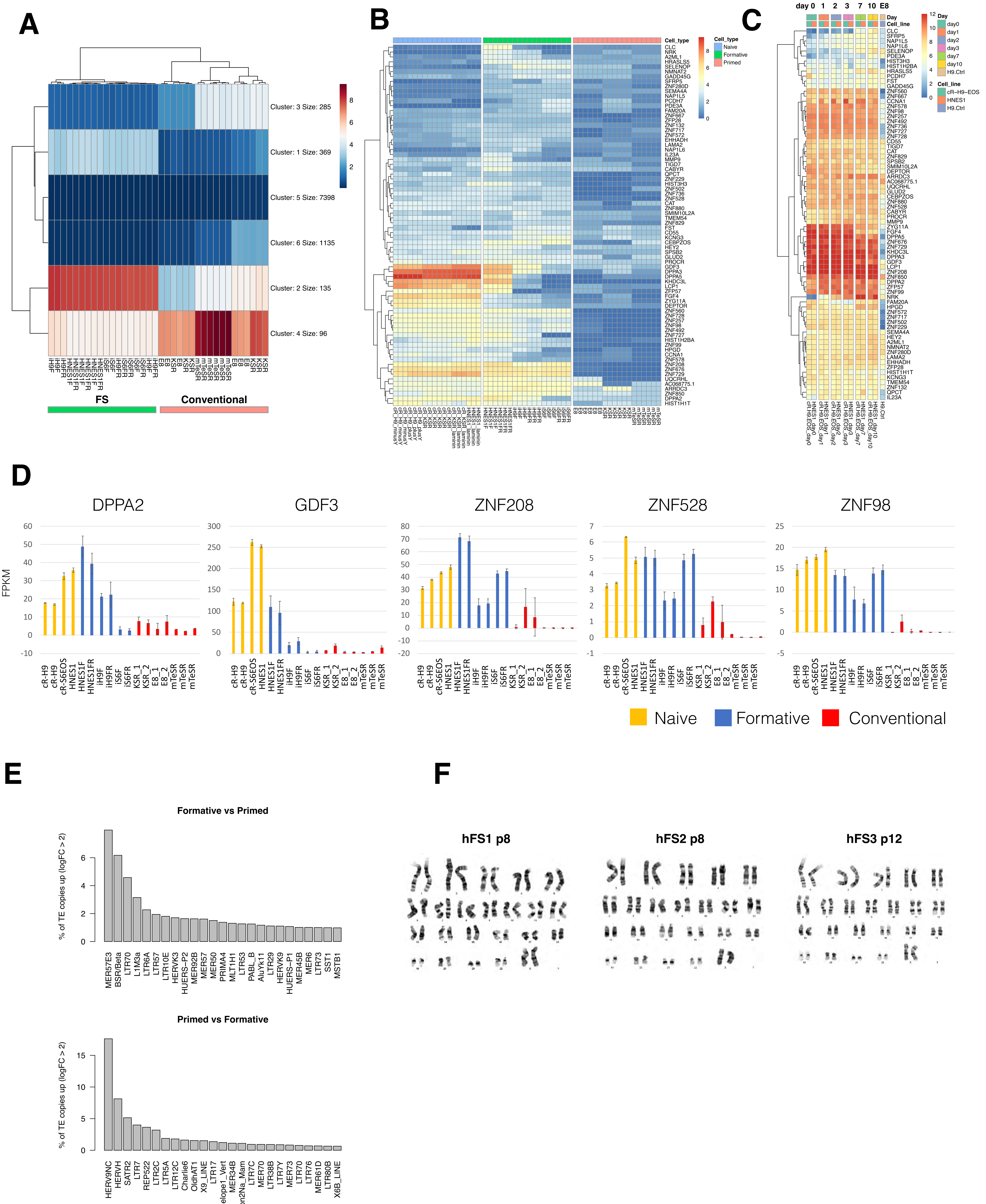

A
Cluster: 3 Size: 285
Cluster: 1 Size: 369
Cluster: 5 Size: 7398
Cluster: 6 Size: 1135
Cluster: 2 Size: 135
Cluster: 4 Size: 96
FS
Conventional
B
Cell_type
Naive
Formative
Primed
C
day 0 1 2 3 7 10 E8
Day
Cell_line
day0
day1
day2
day3
day7
day10
H9.Ctrl
cR-H9-EOS
HNES1
H9.Ctrl
D
DPPA2
GDF3
ZNF208
ZNF528
ZNF98
FPKM
Naive
Formative
Conventional
E
Formative vs Primed
% of TE copies up (logFC > 2)
MER57E3
BSR/Beta
LTR70
L1M3a
LTR6A
LTR57
LTR10E
HERVK3
HUERS-P2
MER92B
MER57
MER50
PRIMA4
MLT1H1
LTR53
PABL_B
AluYk11
LTR29
HERVK9
HUERS-P1
MER45B
MER6
LTR73
SST1
MSTB1
Primed vs Formative
% of TE copies up (logFC > 2)
HERV9NC
HERVH
SATR2
LTR7
REP522
LTR2C
LTR5A
LTR12C
Charlie6
OldhAT1
X9_LINE
LTR17
Penelope1_Vert
MER34B
Helitron2Na_Mam
LTR7C
MER70
LTR38B
LTR7Y
MER73
LTR70
LTR76
MER61D
LTR80B
X6B_LINE
F
hFS1 p8
hFS2 p8
hFS3 p12
